# Biased stochastic motor dynamics drive bidirectional, processive translocation by the AAA+ disaggregase Hsp104

**DOI:** 10.64898/2026.08.09.743785

**Authors:** JiaBei Lin, Jaskamaljot Kaur Banwait, Ayush Saurabh, Kim A. Sharp, Daniel R. Southworth, Steve Pressé, Aaron L. Lucius, Yale E. Goldman, James Shorter

**Affiliations:** Department of Biochemistry and Biophysics, Perelman School of Medicine, University of Pennsylvania, Philadelphia, PA 19104. U.S.A; Department of Chemistry, University of Alabama at Birmingham, Birmingham, AL 35233. U.S.A; Center for Biological Physics, Arizona State University, Tempe, AZ 85287. U.S.A; Department of Physics, Arizona State University, Tempe, AZ 85287. U.S.A; School of Molecular Sciences, Arizona State University, Tempe, AZ 85287. U.S.A; Department of Biochemistry and Biophysics, University of California San Francisco, San Francisco, CA 94158. U.S.A; Institute for Neurodegenerative Diseases, University of California San Francisco, San Francisco, CA 94158. U.S.A; Pennsylvania Muscle Institute, University of Pennsylvania, Philadelphia, PA 19104. U.S.A; Center for Engineering Mechanobiology, University of Pennsylvania, Philadelphia, PA 19104. U.S.A; Department of Physiology, University of Pennsylvania, Philadelphia, PA 19104. U.S.A

**Author notes:** Department of Chemistry and Biochemistry, Rowan University, Glassboro, NJ 08028, U.S.A. Departments of Pharmacology and Cellular and Molecular Biology, University of California, Davis, CA 95616. U.S.A.

## Abstract

Protein aggregation disrupts proteostasis and drives neurodegeneration. Hsp104 is a hexameric, ring-shaped AAA+ ATPase that dissolves protein aggregates, yet how hexamers translocate and extract polypeptides trapped in mechanically resistant aggregates remains unclear. Using substrates that recapitulate the physical constraints of aggregates, we establish that Hsp104 is a processive, bidirectional translocase that can dynamically switch direction while threading a single polypeptide. On mechanically restrained substrates and prions, Hsp104 hexamers execute biased stochastic transitions among three conformational states at individual interprotomer interfaces: closed, extended, and a previously unobserved hyperextended form. These transitions follow kinetically favored paths rather than a rigid rotary sequence. The resulting biased stochastic stepping, enabled by the conformational plasticity of Hsp104 hexamers, underpins operational adaptability and redefines the functional logic of AAA+ motors.

## Background

The AAA+ (ATPases Associated with diverse Activities) protein superfamily comprises molecular machines that harness ATP hydrolysis to remodel diverse substrates, playing critical roles in cellular homeostasis (*1–4*). This family includes critical protein-remodeling factors like Hsp104, VCP/p97, ClpX, katanin, and Skd3, which typically assemble into hexameric rings that translocate polypeptides through a central pore (*1–5*). Hsp104 stands out with a powerful ability to solubilize proteins trapped in otherwise intractable aggregates, thereby safeguarding proteostasis and exhibiting potential as a therapeutic agent against neurodegenerative diseases (*5–27*). Despite these important activities, the precise mechanisms by which Hsp104 engages, translocates, and unfolds substrates during protein disaggregation, and whether these processes are enabled by sequential or stochastic motor processes, remain poorly understood (*5, 6, 28, 29*).

Hsp104 is composed of an N-terminal domain implicated in substrate recognition, nucleotide-binding domain 1 (NBD1), a middle domain (MD) that autoregulates activity and enables collaboration with Hsp70, NBD2, and a short C-terminal domain (*5, 30*). The pore loops found in each NBD of Hsp104, containing conserved tyrosine residues, are essential for substrate engagement and translocation through the axial channel of the hexameric ring (*5, 6, 31, 32*).

Cryo-EM studies have resolved multiple Hsp104 hexamer conformations, including closed, extended, and open states that are distinguished by progressive widening of the seam between protomers P6 and P1 (fig. S1A) (*28, 33*). When Hsp104 pore loops engage the model polypeptide substrate casein, cryo-EM reconstructions predominantly capture closed and extended states, a distribution that has been interpreted as supporting a sequential, rotary translocation mechanism (fig. S1A-C) (*5, 28*). However, this model, based on static snapshots of Hsp104, has not been tested via real-time analysis of conformational dynamics during active translocation. Curiously, the extended state was not observed in ClpB, the prokaryotic homolog of Hsp104 (*34, 35*), underscoring that these two AAA+ machines differ in fundamental aspects of their conformational and functional repertoires (*6, 36*). Indeed, how ClpB translocates polypeptides remains controversial (*37, 38*). Optical tweezers studies suggest a processive, deterministic mechanism involving discrete translocation steps (*37*), whereas single-molecule FRET measurements indicate a diffusive-like mechanism in which ATP primarily regulates directionality rather than directly driving mechanical strokes (*38*). Notably, neither study employed physiologically relevant aggregated substrates (*37, 38*), leaving open how AAA+ disaggregases extract polypeptides buried within mechanically resistant aggregated structures.

Cryo-EM structures of many AAA+ ATPases have spiral-staircase arrangements of pore-loop residues surrounding substrate polypeptides, which has been interpreted to indicate a conserved hand-over-hand mechanism for translocation (*3, 4*). However, such static structural images do not capture the dynamic processes underlying substrate processing, and similar structures can in principle support divergent translocation mechanisms, leaving critical questions about the kinetics of motor function unresolved (*6, 29, 39, 40*). Detailed hydrogen exchange mass spectrometry analysis of Hsp104 has revealed important insights into the structural dynamics and mechanism of Hsp104 hexamers (*41, 42*), but has not resolved the mode of active substrate translocation by individual hexamers. The remarkable capacity of Hsp104 to engage and disassemble diverse substrates ranging from toxic soluble oligomers, denatured protein aggregates, phase-separated condensates, and amyloid or prion fibrils suggests exceptional mechanistic adaptability (*5, 6, 43–45*). However, precisely how a single Hsp104 hexamer translocates diverse polypeptides trapped in complex, heterogeneous aggregated structures remains poorly defined. Indeed, how any AAA+ disaggregase navigates the mechanical resistance and topological complexity presented by protein aggregates, the physiological targets of disaggregase activity, has not been directly addressed.

Here, we address this critical issue and combine single-molecule and ensemble fluorescence experiments to dissect the functional mechanism of Hsp104. We leverage a fully active, cysteine-light Hsp104 variant optimized for site-specific labeling to perform single-molecule fluorescence resonance energy transfer (smFRET) measurements on surface-tethered substrates that recapitulate the mechanical resistance of polypeptides trapped within aggregates. We establish that Hsp104 functions as a processive, bidirectional translocase that can switch direction while threading a single polypeptide, an unexpected behavior for a ring-shaped motor. This switchable bidirectionality may enable complex navigation trajectories suited to the irregular topology of protein aggregates. To peer directly into the Hsp104 engine during active substrate processing, we tracked fluorescent reporters at the inter-protomer seam to monitor conformational states in real time. We demonstrate that Hsp104 exhibits biased stochastic conformational dynamics during substrate translocation, with individual interprotomer interfaces transitioning among three conformations: closed and extended (fig. S1A), anticipated from cryo-EM structures, and a previously unobserved hyperextended state. These transitions proceed along kinetically favored pathways rather than a rigid rotary sequence (fig. S1B, C), a behavior observed on a mechanically restrained substrate and on prions. Together, these findings define Hsp104 as a conformationally plastic motor that can execute processive, bidirectional translocation to enable disaggregation of diverse aggregated substrates. Our results expand the mechanistic paradigm for AAA+ motors and demonstrate how biased stochastic dynamics support functional adaptability.

## Results

### Hsp104 exhibits continuous and switchable, bidirectional translocation of β-casein

To visualize polypeptide translocation events in real time, we employed single-molecule total internal reflection fluorescence (smTIRF) microscopy (Fig. 1A). Single Cy3 fluorophore-labeled Hsp104 was introduced in solution and allowed to interact with surface-anchored β-casein substrates labeled with a single Cy5 fluorophore (Fig. 1A). Casein, an intrinsically disordered protein with a short N-terminal polyproline II-type helical segment and no native cysteines, is a classic model substrate for dissecting the translocation mechanism of Hsp104 (*10, 28*). To enable fluorescence measurements, casein was engineered to contain a single cysteine immediately N-terminal to an Avi-tag inserted at the C-terminal end. This arrangement allowed for site-specific fluorophore labeling adjacent to the C-terminal biotinylation site (see Methods). The resulting construct was immobilized on the coverslip surface via streptavidin–biotin interaction, positioning the fluorophore near the slide surface (Fig. 1A). This surface anchoring physically constrained the casein substrate to mimic the resistance presented by polypeptides trapped within protein aggregates. If Hsp104 engages the N-terminal region and translocates toward the surface-tethered C-terminus, the distance between Hsp104 and the surface-proximal fluorophore should decrease, increasing FRET efficiency; conversely, translocation away from the surface should reduce FRET (Fig. 1A).

**Fig. 1.**
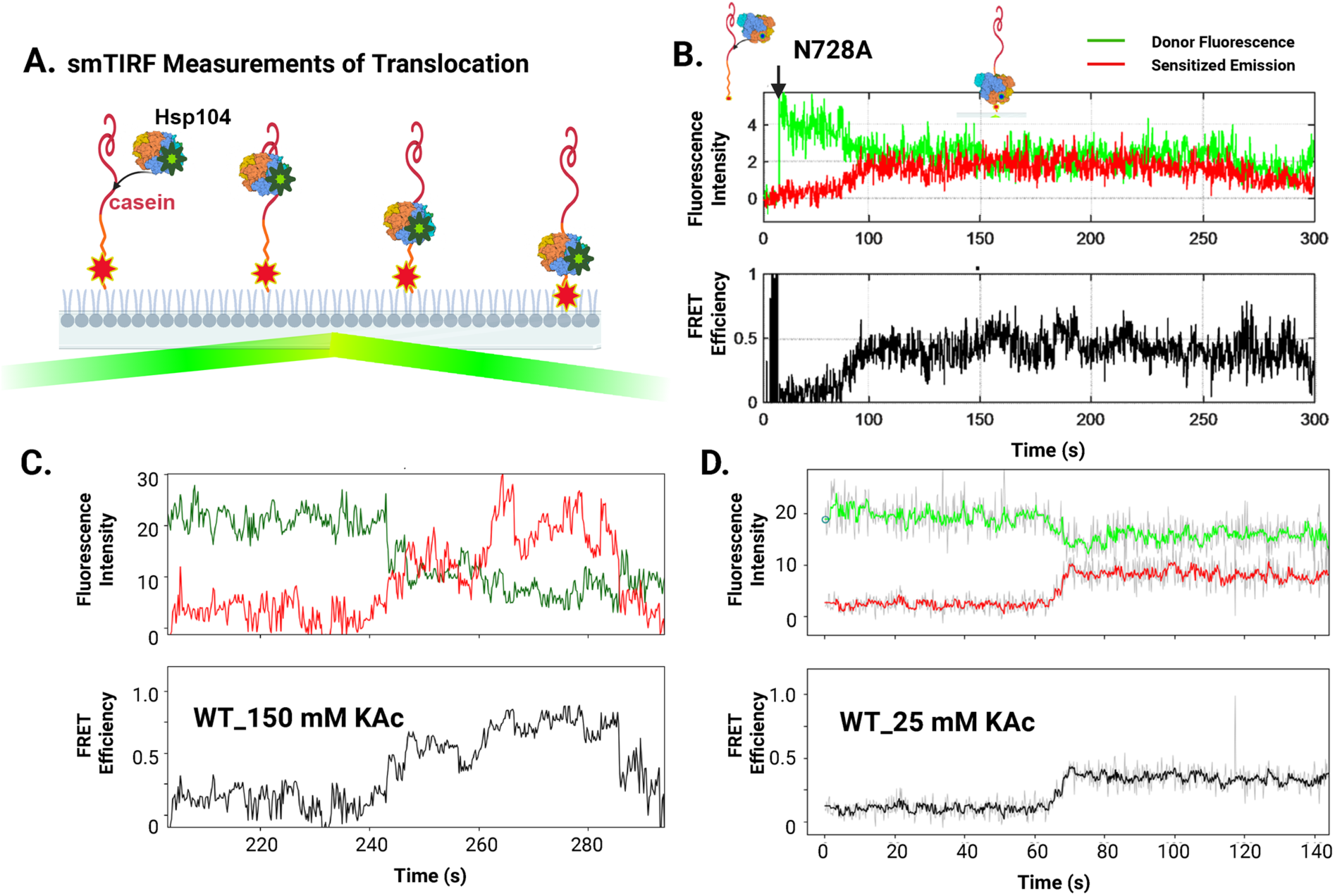
Hsp104 processively translocates along mechanically restrained β-casein. **(A)** Schematic of the smTIRF assay for monitoring Hsp104 translocation. Cy5-labeled (red) β-casein substrates are immobilized on the slide surface via biotin–streptavidin at the C-terminus to mimic the resistance presented by polypeptides trapped within protein aggregates, and Cy3-labeled (green) hexameric Hsp104 binds and translocates along the substrate. **(B)** Representative trajectory of Hsp104^N728A^ moving toward the slide surface on β-casein in HKM150 buffer. Donor (green) and acceptor (red) fluorescence signals are shown unfiltered with the corresponding FRET efficiency trace below. Experiments were performed with 20-60 nM Hsp104^N728A^ (hexameric) and 40-100 nM β-casein loading concentration in 5 mM ATP, using excess β-casein to minimize simultaneous binding of multiple Hsp104 hexamers to the same substrate (see fig. S3). **(C, D)** Representative trajectories for WT Hsp104 translocation on β-casein in HKM150 buffer (C) or HKM25 buffer (D) in the presence of 2.5 mM ATP and 2.5 mM ATPγS. Donor (green) and acceptor (red) fluorescence signals are shown after Chung–Kennedy filtering (*66*), with the calculated FRET efficiency in black. In HKM150 (C), traces were acquired with ≥600 nM total Hsp104 hexamer (including 20 nM Cy3-labeled Hsp104). In HKM25 (D), traces were acquired with 20 nM Hsp104 hexamer following a wash step to remove unbound protein.

For these experiments, we innovated a bespoke single-cysteine Hsp104 variant (MASV-E793C) optimized for site-specific labeling (Supplementary Text and fig. S2A-E). Unless otherwise indicated, all Hsp104 constructs utilized this background, which exhibits similar activity to wild-type (WT) Hsp104 *in vitro* and *in vivo* (fig. S2B, D), and supports the ability of a potentiated Hsp104 variant, Hsp104^E360R^, to mitigate toxicity of aggregation-prone neurodegenerative disease proteins, including α-synuclein, FUS, and TDP-43 (fig. S2E) (*8, 12*).

Hsp104 displays weak affinity for casein in the presence of ATP (*28*). By contrast, an Hsp104 variant bearing an asparagine to alanine mutation in the conserved AAA+ sensor-1 motif in NBD2 (*1*), Hsp104^N728A^, can bind but not hydrolyze ATP at NBD2 and exhibits high-affinity binding to casein in the presence of ATP (*K_D_* ∼20 nM) (*28, 46*). Importantly, Hsp104^N728A^ is an active disaggregase that reactivates luciferase trapped in chemically-denatured aggregates independently of Hsp70 and Hsp40 (fig. S2D) (*47, 48*). Based on these properties, we hypothesized that Hsp104^N728A^ could autonomously translocate substrates at nanomolar concentrations. Similarly, WT Hsp104 reactivates aggregated luciferase in the presence of 1:1 ratio of ATP:ATPγS and in the absence of Hsp70 and Hsp40 (*6, 47, 48*), suggesting that Hsp104 also translocates substrate under these conditions. Both were selected because they support autonomous Hsp104 disaggregase activity at the protein concentrations required for single-molecule experiments, confirming that translocation observed under these regimes reflects a catalytically competent motor (*6, 47, 48*). Thus, we first explored substrate translocation by Hsp104^N728A^ in the presence of ATP (Fig. 1B) and WT Hsp104 in the presence of a 1:1 ratio of ATP:ATPγS (Fig. 1C and D).

To ensure that no more than a single Hsp104^N728A^ hexamer was loaded onto each β-casein polypeptide and to assess potential binding-site preferences, we tested three different loading strategies (Supplementary Text, fig. S3A-C). All three strategies enabled the observation of single Hsp104-Cy3:β-casein-Cy5 complexes distributed within the 200 nm TIRF measurement range (fig. S3A-C). Regardless of the mixing and loading method, in the presence of ATP, we observed intermittent distance changes between Hsp104^N728A^ and the fluorophore-labeled site of β-casein near the coverslip, consistent with substrate translocation (Fig. 1A, B). Strikingly, Hsp104^N728A^ exhibited bidirectional movement along β-casein, with motion both toward (N- to C-terminus) (Fig. 2A) and away (C- to N-terminus) from the slide surface (Fig. 2B). Notably, some smFRET trajectories revealed that Hsp104^N728A^ could move in one direction and then reverse direction (Fig. 2C, D). In ∼9% of cases, Hsp104^N728A^ engaged casein but did not translocate (Fig. 2E, F). Most traces showed bidirectional motion of Hsp104^N728A^ (Fig. 2C, D, F). Among traces with strictly unidirectional motion, translocation was ∼4-fold more frequent from the N- to the C-terminus (Fig. 2F). Overall, these findings reveal that Hsp104^N728A^ can reversibly translocate along a single mechanically restrained polypeptide, demonstrating an unexpected switchable bidirectionality which highlights the adaptability of the motor mechanism. This flexibility challenges the prevailing dogma that Hsp104 simply translocates to one end of the substrate and releases it (*3–5, 28*).

**Fig. 2.**
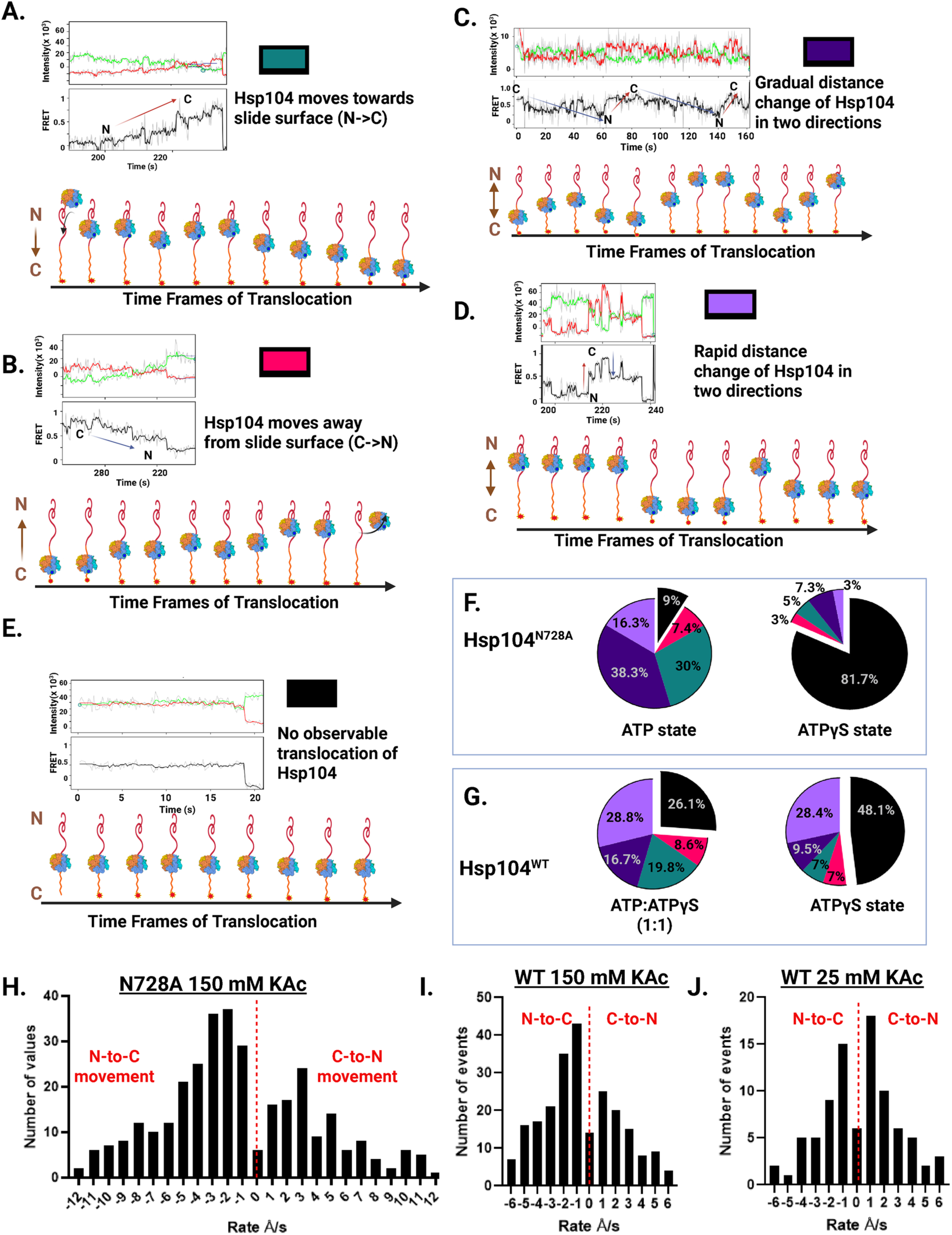
Landscape of Hsp104 translocation trajectories on a single-stranded substrate. **(A–E)** Representative smFRET trajectories illustrating distinct Hsp104^N728A^ translocation behaviors on β-casein. Panels show examples of N-to-C motion toward the slide (A), C-to-N motion away from the slide (B), gradual bidirectional motion (C), rapid bidirectional motion (D), and static binding events (E) for Hsp104^N728A^-Cy3 on surface-anchored β-casein-Cy5, as indicated in the panel labels. The colored square in each trace denotes the trajectory class used in the corresponding pie charts in (F) and (G). **(F)** Classification of Hsp104^N728A^ distance-change events on surface-anchored β-casein in ATP (left) or ATPγS (right), summarizing the fractions of gradual bidirectional (dark purple), rapid bidirectional (light purple), unidirectional N-to-C (green) or C-to-N (pink), and static binding events (black). **(G)** Classification of WT Hsp104 distance-change events on surface-anchored β-casein in ATP:ATPγS (left) or ATPγS (right), using the same color scheme as in (F). **(H–J)** Distributions of apparent translocation rates for Hsp104^N728A^ on surface-anchored β-casein in ATP (H) and for WT Hsp104 on surface-anchored β-casein in ATP:ATPγS under HKM150 (I) and HKM25 (J) buffer conditions. Negative velocities correspond to N-to-C movement toward the slide (increasing FRET), and positive rates correspond to C-to-N movement away from the slide (decreasing FRET); mean values and confidence intervals are reported in Table S1.

Notably, FRET signals often showed progressive increases or decreases in efficiency, consistent with Hsp104^N728A^ gradually translocating along the predominantly unstructured β-casein strand rather than inducing abrupt folding or unfolding events of the casein (Fig. 1B). To assess nucleotide dependence, we repeated the experiments in the presence of ADP or ATPγS, a slowly hydrolyzable ATP analog (Fig. 2F). In ADP, there was no recruitment of Hsp104^N728A^ to casein, consistent with the low affinity of Hsp104^N728A^ for casein in the presence of ADP (*49*). By contrast, in ATPγS, Hsp104^N728A^ bound to β-casein and the complexes were mostly static and exhibited minimal FRET fluctuations (Fig. 2F). Thus, active substrate translocation by Hsp104^N728A^ requires continuous ATP hydrolysis and is effectively halted when hydrolysis is slowed by ATPγS (Fig. 2F).

Occasionally, we observed rapid, reversible distance fluctuations that occurred faster than the camera time resolution (∼200 ms) under both ATP and ATPγS conditions (Fig. 2D, F). These transitions were ∼5.4-fold more frequent in ATP than in ATPγS (Fig. 2F) and likely represent brief bursts of translocation faster than our temporal resolution. An alternative explanation is transient local dynamics within β-casein, but the disordered nature of the substrate makes this possibility less plausible. A further possibility is passive one-dimensional diffusion of Hsp104^N728A^ along the β-casein, in which ATP hydrolysis transiently weakens substrate engagement to permit sliding, whereas ATPγS stabilizes tightly bound states that limit mobility. However, since Hsp104^N728A^ displays similarly high substrate affinity in ATP and ATPγS (*28*), diffusion-based motion seems unlikely.

To determine whether WT Hsp104 exhibits comparable translocase behavior, activity was examined under two buffer conditions that support motor function. Under low-salt conditions (HKM25: 25 mM KAc), which stabilize the hexamer at low concentrations (*46*), WT Hsp104 displayed continuous FRET distance changes in the presence of a 1:1 mixture of ATP and ATPγS, even after unbound protein was removed (Fig. 1D). At more physiological salt concentrations (HKM150: 150 mM KAc), WT Hsp104 likewise translocated along β-casein in both N- to C-terminal (Fig. 1C) and C- to N-terminal directions in the presence of 1:1 ATP:ATPγS (Fig. 2G). Among traces with strictly unidirectional motion, WT Hsp104 moved ∼2.3-fold more frequently from N- to C-terminus than C- to N-terminus (Fig. 2G), indicating reduced directional bias compared to Hsp104^N728A^ (Fig. 2F, G). WT Hsp104 could also reverse direction on a single β-casein strand, and bidirectional traces were most common (Fig. 2G).

In the presence of ATPγS alone under physiological salt conditions, translocation events were less frequent, and all categories of movement, including directional reversals, were reduced (Fig. 2G), indicating that rapid ATP hydrolysis by WT Hsp104 drives efficient substrate translocation. Curiously, rapid, reversible distance fluctuations occurred at similar frequency in 1:1 ATP:ATPγS and ATPγS alone (Fig. 2G), suggesting that these events may arise from sporadic ATPγS hydrolysis by a subset of Hsp104 hexamers, transient β-casein dynamics, or diffusion-like sliding of Hsp104 along the substrate. Together, these findings define Hsp104 as a bidirectional translocase capable of reversing direction on a single substrate.

### Quantifying translocation dynamics of Hsp104^N728A^ and Hsp104

To probe the kinetics of substrate translocation, net distance changes in smFRET trajectories were quantified (fig. S4A), and histograms of average translocation rates were generated (Fig. 2H-J). FRET efficiencies were converted to estimate the distance between the Hsp104 ring and the Cy5 probe near the slide surface (see Methods), with negative distance changes corresponding to N- to C-terminal motion toward the surface (Fig. 2A). Across all events, including both unidirectional and bidirectional traces, Hsp104^N728A^ translocated at an average speed of ∼4.3 Å/s, corresponding to ∼1.26 amino acids per second considering a 3.4 Å contour length per residue along the β-casein chain (*50*) (Fig. 2H; Table S1). This apparent overall rate most likely reflects a mixture of translocation with contributions from local β-casein conformational changes and occasional slipping events (fig. S4A, B). Occasional stepwise distance changes likely represent discrete translocation steps (fig. S4B, pink ovals). Rapid transitions following pauses, which may reflect local unfolding, refolding, slipping, or translocation events faster than our temporal resolution (fig. S4B, red boxes), were excluded from velocity analysis.

WT Hsp104 translocation rates were also quantified. In HKM25 and HKM150 with a 1:1 mixture of ATP and ATPγS, average rates were ∼2.2 Å/s (∼0.65 aa/s; Table S1) for both buffer conditions, which is approximately half the rate of Hsp104^N728A^ in ATP (Fig. 2I, J, Table S1). This ∼2-fold slower translocation rate of WT Hsp104 in 1:1 ATP:ATPγS compared with Hsp104^N728A^ in ATP matches the ∼2-fold lower disaggregase activity of WT Hsp104 under these conditions (*48*). Overall, Hsp104 displays switchable, bidirectional translocation along single-stranded substrates at rates of ∼0.7–1.3 aa/s, underscoring the dynamic and flexible nature of this AAA+ disaggregase.

### Stopped-flow analysis of Hsp104 translocation on structured substrates

To complement the single-molecule findings, ensemble stopped-flow experiments were performed using RepA-Titin substrates containing one to three tandem repeats of the Titin I27 immunoglobulin domain, labeled on a single C-terminal cysteine with Alexa Fluor 555 (AF555; Fig. 3A-a, b) (*51, 52*). The N-terminal 70-residue RepA segment provides a recognition site for Hsp104 (*47*). Unlike β-casein, which is largely unstructured, RepA–Titin substrates present folded domains that require mechanical unfolding during translocation, enabling analysis of how Hsp104 engages more structured substrates. If Hsp104 processively unfolds the Titin I27 domains and translocates from N- to C-terminus, then arrival of Hsp104 at the AF555 probe is expected to produce an increase in fluorescence intensity due to protein-induced fluorescence enhancement (PIFE) (*51, 52*).

**Fig. 3.**
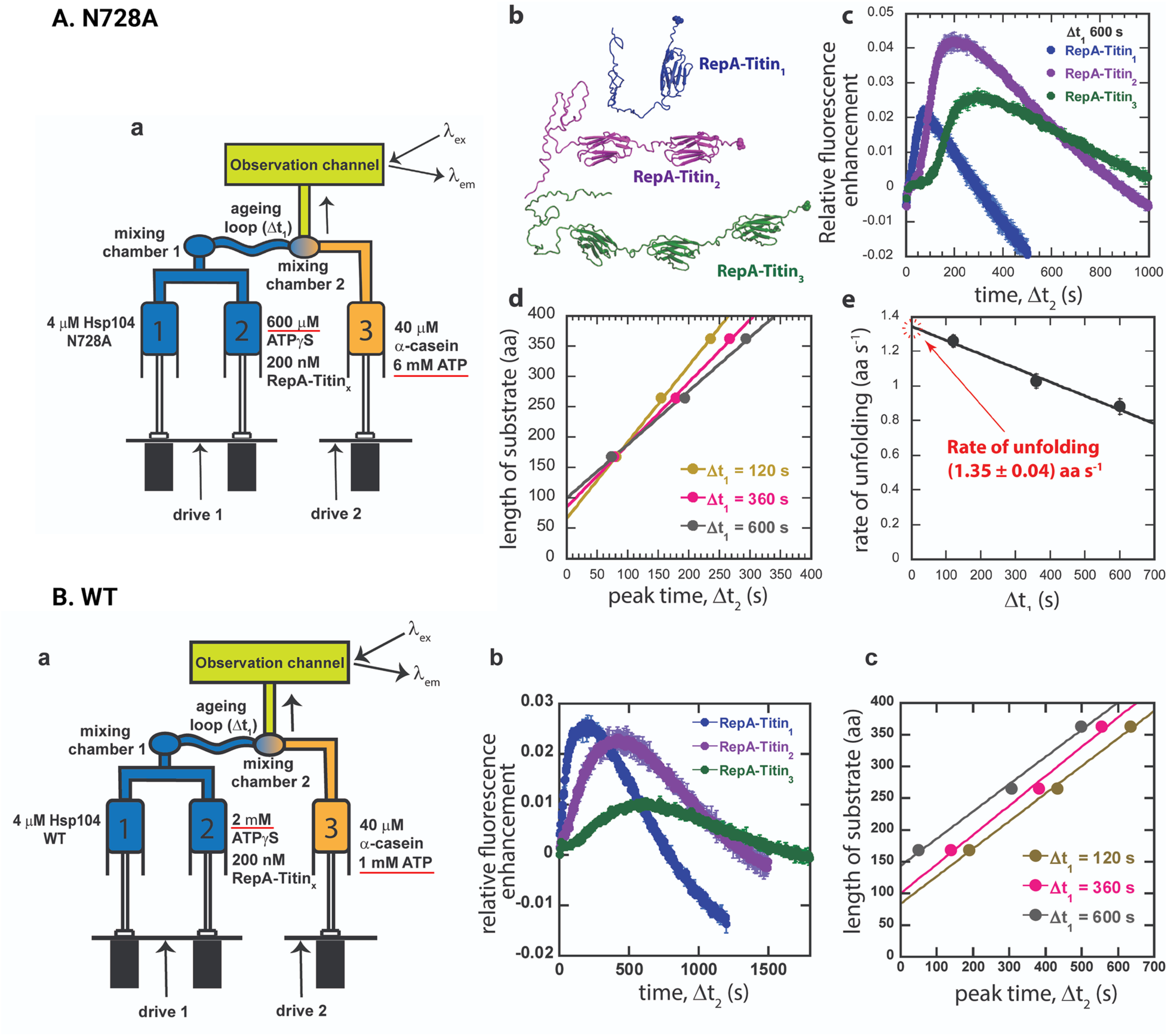
Hsp104^N728A^ and Hsp104 are processive translocases that unfold structured substrates. **(A)** Stopped-flow analysis of Hsp104^N728A^ translocation and unfolding on RepA–Titin_n_ substrates. **(a)** Schematic of the three-syringe, two-mix protocol used to measure single-turnover unfolding and translocation. After an initial aging step in ATPγS to allow hexamer assembly on RepA–Titin_n_ (n = 1, 2, or 3), the mixture is rapidly combined with ATP and an excess of α-casein trap, yielding final concentrations of 1 µM Hsp104^N728A^, 50 nM RepA–Titin_n_, 150 µM ATPγS, 3 mM ATP, and 20 µM α-casein in the observation channel. AF555 fluorescence was excited at 520 nm and emission was collected at >570 nm to report arrival and departure of Hsp104^N728A^ at the C-terminal fluorophore. **(b)** Cartoon of RepA–Titin constructs with one, two, or three Titin I27 domains fused to the RepA recognition module and labeled with AF555 at the C-terminus. **(c)** Representative fluorescence time courses for RepA–Titin_1_ (blue), RepA–Titin_2_ (purple), and RepA–Titin_3_ (green) at Δt_1_ = 600 s. **(d)** Peak fluorescence time as a function of total substrate length for Δt_1_ = 120, 360, and 600 s, with linear fits used to extract apparent unfolding/translocation rates. **(e)** Dependence of the fitted slopes on Δt_1_; extrapolation to Δt_1_ = 0 yields an Hsp104^N728A^ unfolding/translocation rate of ∼1.35 aa s^−1^. **(B)** Stopped-flow analysis of WT Hsp104 translocation and unfolding on RepA–Titin_n_ substrates. **(a)** Schematic of the stopped-flow setup and mixing scheme used to measure WT Hsp104 under single-turnover conditions in a 1:1 mixture of ATP and ATPγS, with α-casein serving as a trap to prevent rebinding. Final concentrations after both mixes are 1 µM Hsp104, 50 nM RepA–Titin_n_, 500 µM ATPγS, 500 µM ATP, and 20 µM α-casein. AF555 fluorescence was monitored as in (A). **(b)** Representative fluorescence time courses for RepA–Titin_1_ (blue), RepA–Titin_2_ (purple), and RepA–Titin_3_ (green) at Δt_1_ = 120 s, averaged from ≥3 sequential traces. **(c)** Peak fluorescence time as a function of total substrate length for Δt₁ = 120, 360, and 600 s; linear fits at each Δt₁ yielded consistent slopes, giving a Δt₁-independent WT Hsp104 unfolding/translocation rate of ∼0.44 aa s⁻¹.

For these experiments, Hsp104 variants containing the native cysteine residues were used rather than the MASV-E793C construct. In one assay configuration, Hsp104^N728A^ (4 µM) in syringe 1 was mixed with 600 µM ATPγS and 200 nM RepA–Titin in syringe 2 (Fig. 3A, panel a). After a defined delay (Δt₁), this aged mixture was rapidly combined with 6 mM ATP and 40 µM α-casein from syringe 3, and fluorescence was then monitored over time (Δt₂). α-Casein served as a competitive trap that prevented rebinding of Hsp104^N728A^ to free RepA-Titin, so the fluorescence signal reported on a single round of unfolding and translocation (*51, 52*). Each mixing step introduced a one-to-one dilution, yielding final concentrations of 1 µM Hsp104^N728A^, 150 µM ATPγS, 50 nM RepA-Titin_n_, 3 mM ATP, and 20 µM α-casein after both mixing events.

Fluorescence time courses displayed characteristic lag, rise, and decay phases (Fig. 3A, panel c) (*51, 52*). The lag phase reflects the time required for Hsp104^N728A^ to unfold and translocate along the substrate before reaching the C-terminus (*51, 52*). The rise phase marks arrival of Hsp104^N728A^ at the AF555-labeled C-terminal end, triggering PIFE (*51, 52*). The decay phase indicates movement of Hsp104^N728A^ away from the fluorophore and likely reflects substrate release averaged with the backtracking observed in the single-molecule experiments (Fig. 2F) (*51, 52*). The time required to reach peak fluorescence increased with substrate length, and a linear fit of peak time versus substrate length yielded a translocation/unfolding rate of ∼1.35 aa/s at Δt₁ = 0 (intercept), consistent with the smFRET rate on β-casein (∼1.26 aa/s) (Fig. 3A, panels d, e; Table S1). Thus, Hsp104^N728A^ engages in processive translocation and unfolding of structured substrates. The similarity of these rates indicates that translocation, rather than substrate unfolding, is rate limiting: Hsp104^N728A^ is not detectably impeded by unfolding the stably folded β-sandwich Titin I27 domain. In this way, Hsp104^N728A^ behaves more like a constant-velocity winch than a cautious, obstacle-sensing motor, pulling substrates through at a similar rate whether confronted with unfolded β-casein or stably folded Titin I27 domains.

To assess WT Hsp104, the stopped-flow assay was repeated using a 1:1 mixture of ATP and ATPγS (Fig. 3B). Syringe 1 contained 4 µM Hsp104, and syringe 2 contained 200 nM RepA–Titin and 2 mM ATPγS. After a defined preincubation (Δt₁ = 120, 360, or 600 s), the aged sample was rapidly mixed with 1 mM ATP and 40 µM α-casein trap (Fig. 3B, panel a), yielding final concentrations of 1 µM Hsp104, 500 µM ATPγS, 50 nM RepA–Titin_n_, 500 µM ATP, and 20 µM α-casein after both mixing events.

For each Δt₁, WT Hsp104 produced reproducible PIFE traces, with peak times that increased with substrate length (Fig. 3B, panel b). Linear fits of substrate length versus time of peak fluorescence gave a translocation/unfolding rate of ∼0.44 aa/s, independent of Δt₁ (Fig. 3B, panel c; Table S1). This rate is ∼3-fold slower than that of Hsp104^N728A^ and broadly comparable to, albeit slightly lower than, the value obtained from smFRET analysis of WT Hsp104 on β-casein (∼0.65 aa/s), consistent with the same motor operating within a similar rate regime across substrates and assay platforms. These findings are consistent with the elevated disaggregase activity of Hsp104^N728A^ in ATP compared with WT Hsp104 in 1:1 ATP:ATPγS (*48*). Thus, WT Hsp104, like Hsp104^N728A^, processively unfolds and translocates structured substrates, but under these conditions operates more slowly, behaving more like a lower-gear version of the same motor that pulls steadily through Titin I27 domains at a reduced yet still processive pace.

### Hsp104^N728A^ hexamers exhibit heterogeneous conformational dynamics at individual interprotomer interfaces, with three distinct states

Having established that Hsp104 is a processive translocase, we next dissected the internal subunit interface movements that power translocation. Individual Hsp104^N728A^ MASV–E793C subunits were labeled with Cy3 or Cy5 at residue 793, and hexamers were assembled by mixing Cy3-Hsp104^N728A^, Cy5-Hsp104^N728A^, and unlabeled Hsp104^N728A^ at 1:1:4 or 2:2:2 ratios to produce hetero-hexamers containing both labeled and unlabeled subunits within a single assembly (Fig. 4A) (*6*). Residue 793 was selected because cryo-EM structures predict substantial inter-subunit distance changes as the spiral seam between protomers P6 and P1 widens and narrows (Table S2) (*5, 28*). Fluorophores at C793 in adjacent non-seam subunits in closed and extended state hexamers are expected to show uniformly high FRET (∼0.70–0.77), whereas the P6–P1 seam interface is conformationally diagnostic, with high FRET in the closed state (0.64; PDB: 5VJH), intermediate FRET in the extended state (0.41; PDB: 5VYA), and negligible FRET in open states (fig. S1A–C, Fig. 4A, fig. S2C, Table S2) (*5, 28, 33*). Labeled subunits separated by one or more intervening protomers (≥10 nm apart) are expected to produce no measurable FRET (fig. S1A, Fig. 4A, fig. S2C) (*5, 28, 33*). Together, this design enables direct monitoring of how frequently the Hsp104 seam transitions between closed, extended, and open conformations during translocation, and whether these transitions follow a defined sequence (fig. S1A–C, Fig. 4A).

**Fig. 4.**
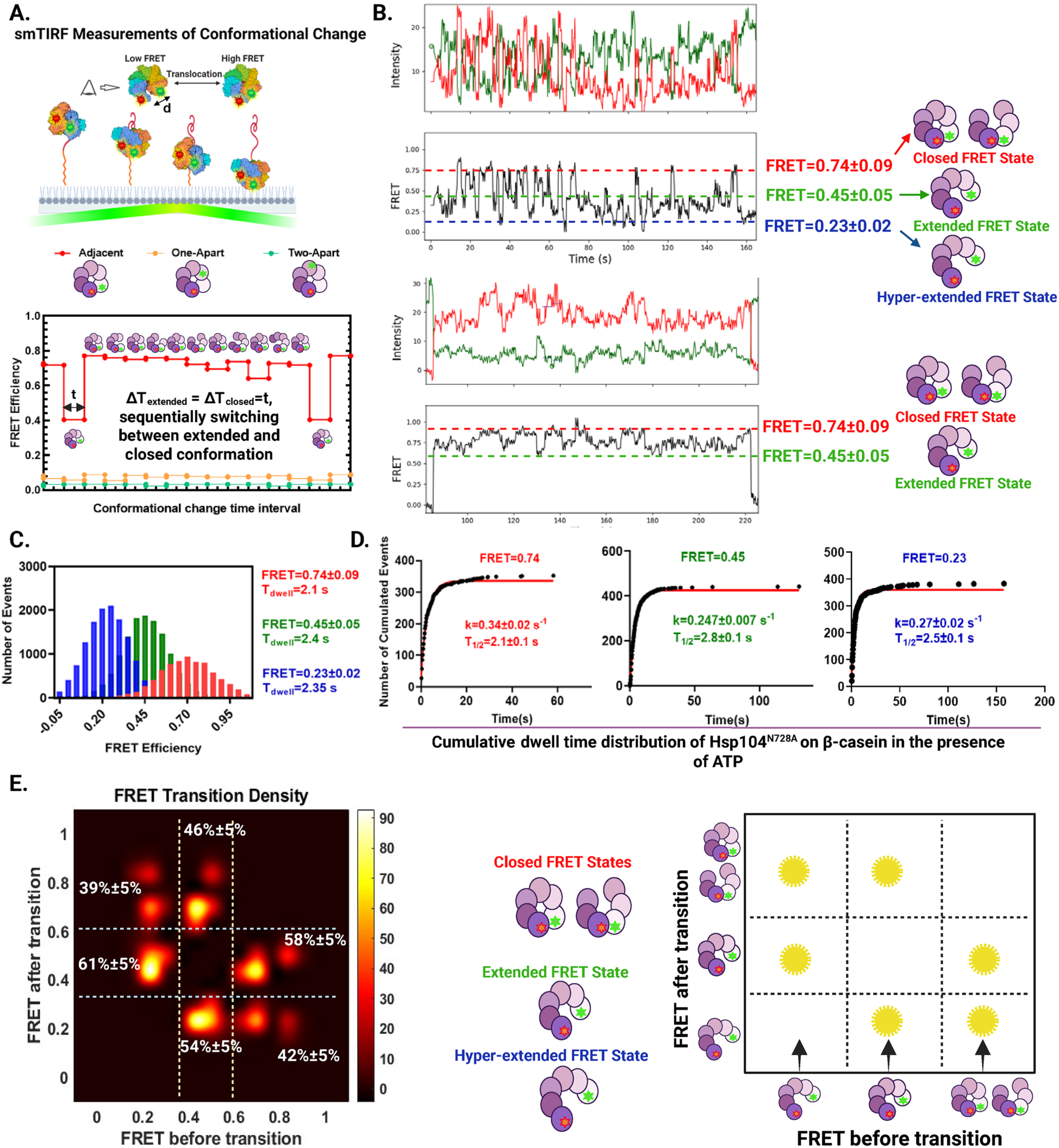
Hsp104^N728A^ exhibits three-state conformational dynamics and operates as a biased stochastic motor. **(A)** Schematic of the smFRET assay for monitoring Hsp104 conformational changes during translocation. Heterohexamers were assembled from mixtures of Cy3-labeled, Cy5-labeled, and unlabeled Hsp104^N728A^ subunits at defined ratios, and FRET efficiency between labels at residue 793 in adjacent protomers was used to report intersubunit distances. Simulated FRET trajectories illustrate the behavior expected for an ideal sequential rotary mechanism in which the spiral seam moves stepwise around the ring and subunits alternate between closed (PDB: 5VJH) and extended (PDB: 5VYA) conformations. **(B)** Representative smFRET trajectories of Hsp104^N728A^ hexamers on β-casein, showing donor (green), acceptor (red), and calculated FRET efficiency (black) after Chung-Kennedy filtering. Dashed horizontal lines indicate the mean FRET efficiency for each hidden-Markov-model-assigned state, with representative hexamer schematics and state assignments (closed, extended, hyperextended) shown at right. Across the data set (n = 105 hexamers total), the ebFRET-refined trajectory classification used for the subpopulation analysis in fig. S5 assigned 70.5% of Hsp104 hexamers to a three-state model (top panels) and 29.5% to a two-state model (bottom panels); the corresponding per-trace Bayesian mixture assignment (Table S3) gave 63.8% and 36.2%, respectively. **(C)** Global ebFRET analysis of 105 trajectories using a three-state model identified mean FRET efficiencies of ∼0.74, ∼0.45, and ∼0.23, corresponding to closed, extended, and hyperextended conformations, respectively. **(D)** Cumulative dwell-time distributions for each state were fitted with single exponentials to obtain state-specific rate constants and half-times (n = 352, 440, and 383 dwells for closed, extended, and hyperextended states, respectively, see Equation 4). **(E)** Transition density plot (left) summarizing the preferred conformational pathways among the three states, and corresponding schematic (right). Transitions are enriched between neighboring states but also include direct switches between closed and hyperextended conformations, consistent with biased stochastic conformational stepping. The transition-probability estimates and confidence intervals are derived from non-parametric bootstrap resampling of pooled trajectories and describe the empirical kinetic landscape at labeled inter-protomer interfaces.

Cryo-EM studies led to the proposal that Hsp104 translocates substrates via a sequential rotary mechanism (fig. S1A-C) (*4, 28*), but this model, derived from static structural snapshots, has not been tested in real time (*29*). Mutant subunit doping experiments argue against a strictly sequential rotary mechanism for labile substrates (*6*). However, for more stable substrates (for example, amyloids) all six WT subunits are required, leaving open the possibility that a rotary mechanism operates under specific conditions (*6*). A similar sequential rotary scheme has been proposed for many ring-shaped AAA+ translocases based on cryo-EM structures, yet direct visualization of translocation in real time is lacking (*3, 4*). If a rotary model (fig. S1A-C) applies, individual subunits should undergo periodic transitions in a defined order, with the spiral seam cycling regularly across the interface between labeled protomers. To capture this behavior, idealized rotary transitions were simulated assuming equal dwell times in closed and extended states (Fig. 4A), in which the intermediate-FRET state at the seam of the extended state would be visited only briefly, occupying on the order of 1/12 of the full cycle.

Experimental FRET trajectories deviated markedly from the rotary prediction (Fig. 4B). Instead of brief (∼1/12 of total occupancy) visits to the intermediate-FRET state (Fig. 4A), Hsp104^N728A^ subunits frequently remained in extended conformations for prolonged periods (Fig. 4B-D, fig. S5A-C). To classify trajectories, we used ebFRET, an empirical Bayes hidden Markov modeling approach that infers FRET states and transition kinetics from single-molecule time series (*53*). Under the ebFRET-refined trajectory classification used for the subpopulation analysis, 70.5% of Hsp104^N728A^ traces were best described by three discrete FRET states and 29.5% by two states (fig. S5D). The corresponding fractions under the per-trace Bayesian mixture classification (Table S3) were 63.8% (three-state) and 36.2% (two-state); the two methods are semi-independent (see Methods) (*54*) and agree that the three-state category is dominant. In the per-trace Bayesian mixture analysis, one-, four-, and five-state solutions were, in every trace, several orders of magnitude less likely than the two- or three-state descriptions. This method provides per-trace evidence against both simpler and more complex models. As an independent confirmatory analysis, model-order comparison using variational lower bounds from ebFRET (*53*) likewise showed the largest gain in approximate model evidence at the 2→3 state transition, with progressively diminishing returns for additional states (Tables S4, S5). Together, these two independent analyses support a three-state model as the most parsimonious and biologically interpretable description of Hsp104^N728A^ conformational dynamics, with the ∼10^3^-unit gain in mean lower bound at the 2→3 state transition arguing against overfitting to noise or kinetic substructure (see Supplementary Text; Table S4, S5).

The three most probable FRET efficiencies were ∼0.23, ∼0.48, and ∼0.74 by Bayesian mixture model analysis (*54*), whereas ebFRET (*53*) yields the same modal values of ∼0.23, ∼0.45, and ∼0.74 within the error. Notably, the two methods optimize different objective functions but identify the same three physical states. We designate these states as hyperextended (low FRET), extended (intermediate FRET), and closed (high FRET), respectively (Table S3).

The extended and closed states are anticipated from prior cryo-EM structures (fig. S1A–C) (*28*), but the hyperextended state was unanticipated and has not been observed in previous structural studies. Importantly, the hyperextended state is not simply the previously described open states (fig. S1A), which are predicted to produce negligible FRET (Table S2), reflecting complete disengagement of the P6–P1 NBD2–NBD2 interface (*28, 33*). By contrast, the hyperextended state yields low but measurable FRET (∼0.23), indicating that adjacent protomers at the seam remain in partial contact (fig. S1A; Table S2). The hyperextended state is therefore structurally distinct from both the extended and open states.

Importantly, the 29.5% of traces best described by two states in the ebFRET-refined analysis (fig. S5D) and 36.2% in the Bayesian mixture model (Table S3) do not represent a mixing artifact. Subpopulation analysis confirmed that these hexamers, when grouped by their refined two-state assignment, showed internally consistent FRET values, either hyperextended plus extended (fig. S5A) or extended plus closed (fig. S5B), distinct from the pooled three-state distribution (fig. S5C). Thus, three-state traces reflect genuine within-hexamer behavior rather than an average of two-state populations.

The extended and hyperextended states, both reporting a widened P6–P1 seam, together occupied ∼78.3% of the recording period per Hsp104^N728A^ hexamer (Fig. 4C, Table S6), far exceeding the occupancy predicted by a strictly sequential rotary mechanism (Fig. 4A, Table S6). In the simplest formulation, where a single interface occupies the seam for 1 of 12 cycle sub-steps, the widened seam is predicted to be occupied only ∼8% (1/12) of the time (Fig. 4A). However, because our data resolve two distinct widened conformations rather than one, we extended this null model to allow the seam interface to sample both widened sub-states in succession, yielding a more permissive prediction of ∼16.7% (2/12) of the time (Table S6). Even against this generous benchmark, observed widened-state occupancy exceeds the rotary prediction nearly fivefold. Hsp104^N728A^ therefore dwells predominantly in widened seam architectures (i.e., extended and hyperextended), making comparatively brief excursions to the closed state. This pattern is incompatible with the evenly distributed state occupancies of a rigid rotary scheme (Fig. 4A; fig. S1B, C) (*28*).

### Hsp104^N728A^ functions as a biased stochastic motor with preferred transition pathways

Dwell time distributions showed that each of the three conformations persisted for ∼2.1–3.7 seconds on average, indicating substantial stability during substrate processing (Fig. 4C, D; fig. S5A-C). Transition density plots revealed that exchanges between neighboring conformational states occurred more frequently than direct transitions between the hyperextended and closed states (Fig. 4E; Table S7). Quantitatively, transitions from the hyperextended state proceeded most frequently to the neighboring extended state (∼61%) rather than directly to the closed state (∼39%), whereas transitions from the closed state likewise favored the adjacent extended state (∼58%) over the hyperextended state (∼42%). Thus, Hsp104^N728A^ behaves as a biased stochastic motor, exhibiting a preference for neighboring-state exchange within the three-state conformational network while retaining a substantial minority of direct long-range transitions between the hyperextended and closed states.

### WT Hsp104 functions as a biased stochastic motor with preferred transition pathways

To assess whether WT Hsp104 shares the same conformational logic, 149 FRET trajectories from five independent replicates were measured for WT Hsp104 in the presence of a 1:1 mixture of ATP and ATPγS, which supports translocation on β-casein (fig. S6A, B). Using ebFRET, three FRET states were again identified, with mean efficiencies of 0.27 (hyperextended), 0.50 (extended), and 0.69 (closed), and corresponding dwell times of 5.84, 5.66, and 5.2 seconds, respectively (fig. S6A-D; fig. S7A-C; Tables S3, S4, and S5). As with Hsp104^N728A^, most WT hexamers (∼73%) sampled all three conformational states, while a minority (∼27%) showed only two-state behavior (hyperextended plus extended or extended plus closed; fig. S7D). The extended and hyperextended states together occupied ∼71.4% of the recording period per hexamer (fig. S6C, S7A-C), again far exceeding the rotary prediction (Fig. 4A; Table S6).

Cumulative dwell-time distributions showed that the average rate constant for each WT state transition was approximately half that of Hsp104^N728A^ (fig. S6D), in line with the ∼2-fold slower WT translocation rate (Table S1). These findings are consistent with tight coupling between conformational cycling and substrate translocation. Transition density analysis revealed the same qualitative behavior as Hsp104^N728A^: transitions between neighboring states occurred more frequently than direct hyperextended↔closed transitions, with quantitatively similar frequencies (fig. S6E; Table S7). Thus, WT Hsp104 also behaves as a biased stochastic motor, like Hsp104^N728A^, sharing both the three-state conformational architecture and the neighboring-state transition bias when translocating on β-casein.

### Hsp104 exhibits biased stochastic conformational dynamics on Sup35-NM prions in ATP

To investigate Hsp104 dynamics on a natural aggregated substrate and to provide a critical control that eliminates any potential influence of ATPγS or the N728A mutation on conformational dynamics, interactions with Sup35-NM prion fibrils were examined using WT Hsp104 in ATP alone. The N-terminal (N) prion domain of Sup35 forms the amyloid core of infectious fibrils that encode the yeast prion [*PSI*^+^], while the adjacent, highly charged middle (M) domain of Sup35 modulates fibril assembly and stability; together, these two domains (NM) are commonly used as a minimal prion-forming fragment (*55–59*). Sup35-NM prion fibrils are strongly engaged and disassembled by WT Hsp104 in the presence of ATP, whereas Hsp104^N728A^ is unable to dissolve these structures (*6, 43, 57, 60–62*). Sup35-NM fibrils were prepared by gentle agitation overnight (*57, 60*). As with β-casein, WT Hsp104 displayed heterogeneous FRET dynamics in the presence of Sup35-NM fibrils (Fig. 5A, B).

**Fig. 5.**
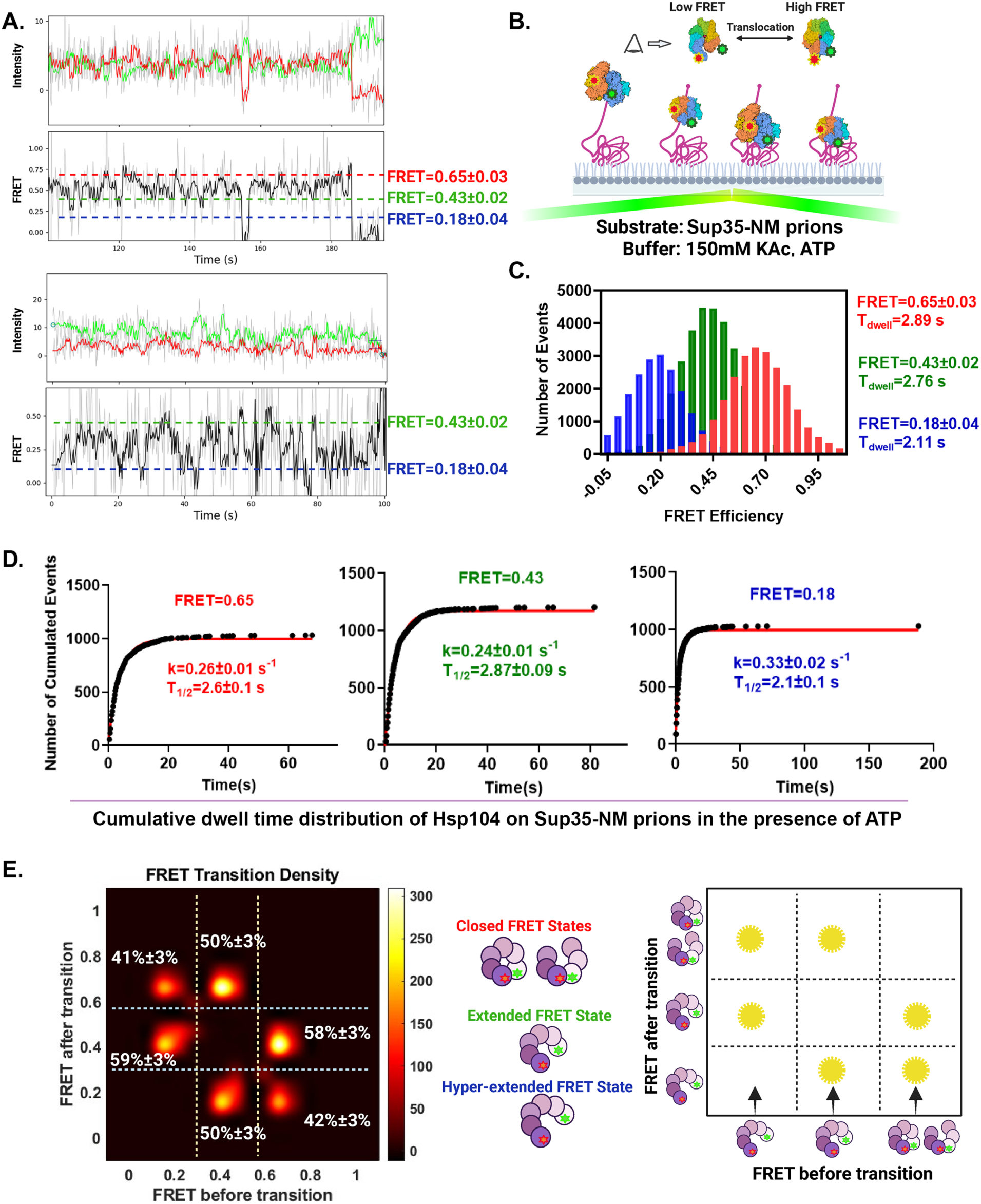
WT Hsp104 exhibits three-state, biased stochastic conformational dynamics on Sup35-NM prions. **(A)** Conformational dynamics of WT Hsp104 on Sup35-NM fibrils in ATP. Representative smFRET trajectories of WT Hsp104 hexamers bound to Sup35-NM fibrils in HKM150 buffer with 5 mM ATP, showing donor (green), acceptor (red), and FRET efficiency (black) with Hidden-Markov state optimization. Dashed horizontal lines indicate the mean FRET efficiency for each assigned state (closed, 0.65±0.03; extended, 0.43±0.02; hyperextended, 0.18±0.04). Across the data set (n = 169 hexamers from six independent experiments), the ebFRET-refined trajectory classification used for the subpopulation analysis in fig. S8 assigned 84.6% of Hsp104 hexamers to a three-state model and 15.4% to a two-state model; the corresponding per-trace Bayesian mixture assignment (Table S3) gave 80.5% (three-state) and 19.5% (two-state). **(B)** Schematic of the experimental setup for measuring WT Hsp104 conformational changes on Sup35-NM fibrils. **(C)** Global ebFRET analysis of the 169 hexamers using a three-state model identified mean FRET efficiencies of ∼0.65, ∼0.43, and ∼0.18, corresponding to closed, extended, and hyperextended conformations, with mode dwell times of 2.89, 2.76, and 2.11 s, respectively. **(D)** Cumulative dwell-time distributions for each state were fitted with single exponentials to obtain state-specific rate constants and half-times (n = 1033, 1203, and 1030 dwells for closed, extended, and hyperextended states, respectively, see Equation 4). **(E)** Transition density plot (left) summarizing the preferred conformational pathways among the three states, and corresponding schematic (right). Transitions are enriched between neighboring states but also include direct switches between closed and hyperextended conformations, consistent with biased stochastic conformational stepping. The transition-probability estimates and confidence intervals are derived from non-parametric bootstrap resampling of pooled trajectories and describe the empirical kinetic landscape at labeled inter-protomer interfaces.

ebFRET analysis of 169 trajectories from six independent experiments identified three FRET states with mean efficiencies of 0.18 (hyperextended), 0.43 (extended), and 0.65 (closed), and corresponding dwell times of 2.11, 2.76, and 2.89 seconds, respectively (Fig. 5A, C, D; fig. S8A--C; Tables S3, S4, and S5). These dwell times are similar to those observed for Hsp104^N728A^ on β-casein in ATP and approximately half those measured for WT Hsp104 on β-casein in 1:1 ATP:ATPγS (Fig. 5C, D), indicating that both substrate and nucleotide context modulate conformational kinetics. Against Sup35-NM prions, more Hsp104 hexamers (∼85%) sampled all three conformational states, whereas a minority showed two-state behavior, and hexamers that sampled only extended plus closed states were rare (∼1.8%; fig. S8D). Combined hyperextended plus extended occupancy in WT Hsp104 on Sup35-NM prions was ∼68.1% (Fig. 5C, Table S6), exceeding the rotary prediction and confirming that predominant widened-seam occupancy is preserved on a prion substrate. Transition density analysis again revealed predominant transitions between neighboring conformational states with less frequent direct transitions between the hyperextended and closed conformations (Fig. 5E; Table S7). Thus, WT Hsp104 also behaves as a biased stochastic motor on the natural prion substrate Sup35-NM, preserving the same three-state conformational architecture and neighboring-state transition bias during prion remodeling.

### Hsp104 cycling is stochastic but kinetically biased

To further test whether conformational cycling of Hsp104 follows a rigid, periodic trajectory or a more flexible, stochastic scheme across all three experimental regimes, a model-independent autocorrelation function (ACF) analysis was applied to smFRET trajectories (*63*) (see Supplementary Text; figs. S9, S10). Periodic processes produce peaks in the ACF at delay times matching the underlying period, but such peaks were not observed in any of the conditions examined, arguing against strictly periodic behavior, and strongly disfavoring a sequential rotary mechanism (see Supplementary Text; figs. S9, S10). Nevertheless, certain conformational transitions occurred more frequently than others, indicating that stochastic behavior is biased by kinetic preferences toward preferred pathways (see Supplementary Text; figs. S9, S10).

### Hsp104 maintains similar transition architecture across substrates

Finally, we compared conformational exchange behavior among all three experimental regimes. Across all conditions, transitions between neighboring conformational states were consistently more frequent than direct transitions between the hyperextended and closed conformations. For example, transitions from the hyperextended state to the neighboring extended state occurred at frequencies of ∼61%, ∼64%, and ∼59% for Hsp104^N728A^–β-casein, WT Hsp104–β-casein, and WT Hsp104–Sup35-NM prions, respectively, whereas direct hyperextended to closed transitions occurred at lower frequencies of ∼39%, ∼36%, and ∼41% (Fig. 4E, Fig. 5E, fig. S6E; Table S7). Similarly, transitions from the closed state favored exchange with the neighboring extended state (∼58–68%) over direct transitions to the hyperextended state (∼32–42%) across all three conditions. Thus, the motor does not appear to markedly rewire its transition logic depending on the substrate. Instead, the same three-state stepping rules are maintained across conditions, with only modest quantitative differences in the relative abundance of some long-range transitions, particularly during remodeling of Sup35-NM fibrils. Overall, these results indicate that substrate and nucleotide context modulate how often each state is occupied and how long it persists, but not the fundamental transition architecture. Hsp104 retains a conserved three-state conformational framework with preferential neighboring-state exchange during active substrate processing, consistent with non-periodic conformational cycling rather than rigid sequential rotary stepping.

## Discussion

Using single-molecule and ensemble approaches, our study redefines Hsp104 as a conformationally plastic, processive motor that translocates substrates via biased stochastic dynamics (Fig. 6). We establish that Hsp104 exhibits unexpected, switchable bidirectional movement on mechanically restrained polypeptides, with individual interprotomer interfaces transitioning among three distinct conformations: closed, extended, and a previously undetected hyperextended state, without following a rigidly fixed transition sequence or successive rotary widening of the interprotomer seam. Instead, the motor samples these states along kinetically favored, non-sequential pathways, a behavior that may enable adaptive navigation and solubilization of the complex and irregular topologies of diverse protein aggregates.

**Fig. 6.**
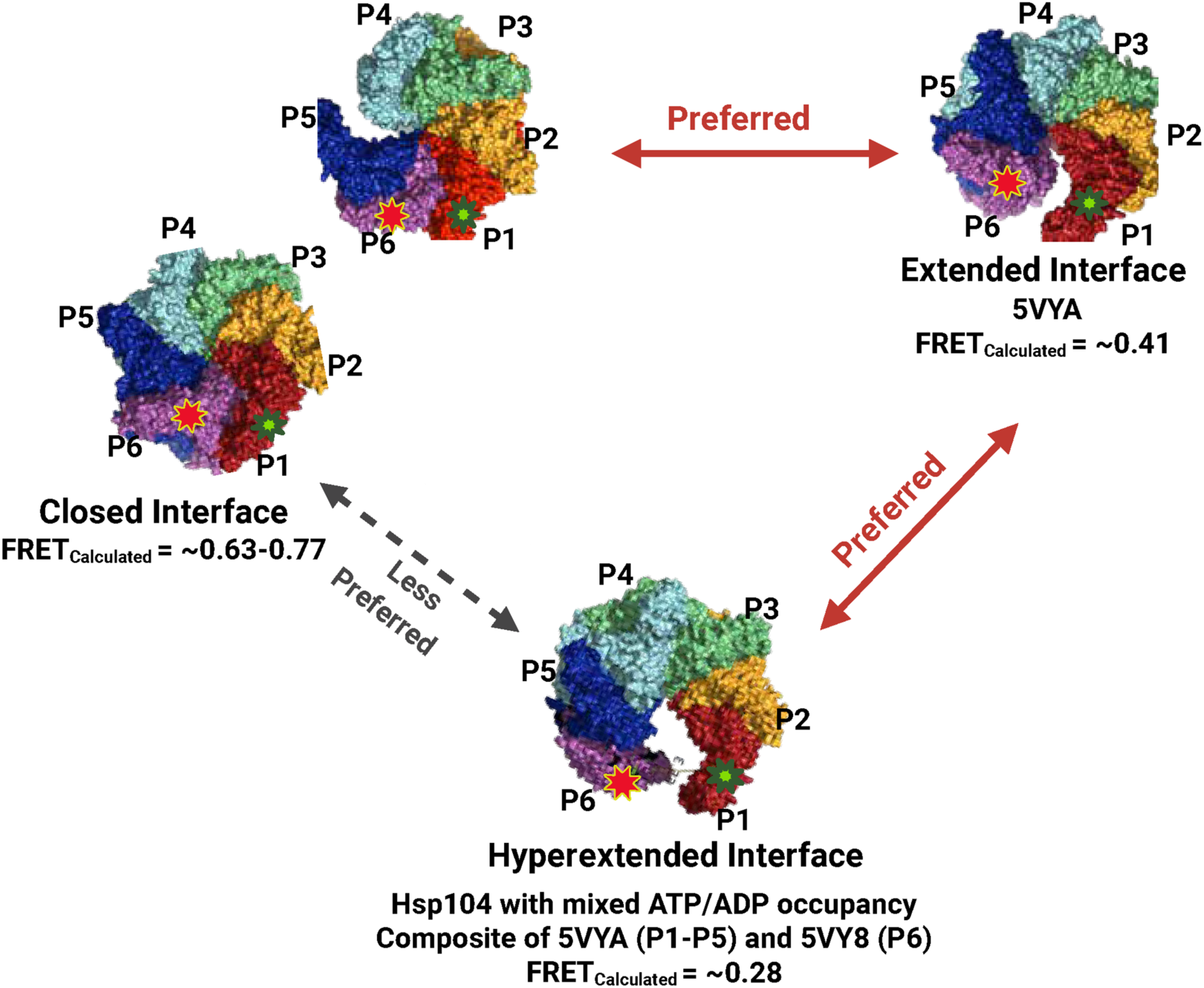
Hsp104 operates as a biased stochastic motor. Each interprotomer interface within the Hsp104 hexamer can adopt one of three conformations — closed, extended, or hyperextended — with preferred transitions between them during substrate translocation. Stars denote an example in which fluorophores at residue 793 are found on protomers P1 and P6, reporting the conformation of that interface by interprotomer FRET. A closed interface (represented by PDB: 5VJH and 5VYA, both ATPγS-bound; closed interfaces can be found in any nucleotide-bound state, see Table S2) exhibits high FRET (FRET_calculated_ ∼0.63–0.77; Table S2); an extended interface (PDB: 5VYA, ATPγS-bound) exhibits intermediate FRET (FRET_calculated_ ∼0.41); and a hyperextended interface exhibits low FRET (FRET_calculated_ ∼0.28). The hyperextended conformation is modeled as a composite of 5VYA (protomers P1–P5) docked with 5VY8 (protomer P6, ADP-bound open state), producing a wider seam opening than captured in either parent structure. Transition probabilities indicate that interfaces preferentially move between adjacent conformations, i.e., closed↔extended and extended↔hyperextended, but can also switch directly between the more distant closed and hyperextended conformations, consistent with a biased stochastic stepping mechanism. This three-conformation model contrasts with sequential, rigid-geometry motors (fig. S1), revealing how Hsp104 achieves processive translocation through conformationally flexible, non-periodic dynamics.

This switchable, probabilistic operation, observed directly in real time on an active motor rather than inferred from static snapshots, marks a clear departure from the deterministic rotary model long proposed for AAA+ motors (*3, 4*). Cryo-EM reconstructions have been interpreted to support a rigid, hand-over-hand mechanism (*3–5, 28*), yet the real-time smFRET data reveal no periodicity and show that the extended state is occupied far longer than the 1/12 or 1/6 average expected for the rotary scheme. Prior structural studies captured the Hsp104 ring in defined conformational poses (*28, 33*), but by tracking fluorescent reporters at the inter-protomer seam in real time, we could watch that seam widen and narrow as Hsp104 engaged and processed substrate, something no structural snapshot can reveal. Across all three experimental conditions, hexamers spent ∼68–78% of their recording time in extended or hyperextended conformations, far exceeding the ∼8% occupancy expected for a simple rotary cycle (Fig. 4A) and even the ∼16.7% expected if both widened-seam conformations are included (Table S6).

Rather than distributing time evenly across conformations as a rotary motor would, Hsp104 traverses a flexible energy landscape in which transitions between conformationally adjacent states are strongly preferred, yet direct jumps between the more distant hyperextended and closed states can also occur. These long-range transitions bypass the extended state and may reflect larger conformational excursions that contribute to processivity on structurally heterogeneous aggregates. Modest differences in the relative abundance of long-range transitions were observed across conditions, particularly during remodeling of Sup35-NM prion fibrils, but these should be interpreted cautiously given that inferred transition frequencies depend on model-derived state assignments and finite temporal resolution. Taken together, our findings support a conserved but non-rigid conformational framework. This flexible conformational sampling may facilitate rapid direction reversals, expanding the range of trajectories available to the motor and enabling Hsp104 to escape dead ends or mechanically constrained regions within aggregated proteins. Such adaptability may drive efficient polypeptide extraction from the diverse aggregated assemblies encountered in cells (*6, 44*).

Beyond conformational plasticity, the stopped-flow translocation assays reveal a striking mechanistic distinction between Hsp104 and its bacterial homolog ClpB. For ClpB, unfolding is rate-limiting: the motor pauses at each folded Titin I27 domain and cooperatively unfolds all ∼98 amino acids before rapidly translocating the nascent unfolded chain (*51, 52*). By contrast, Hsp104 is largely unimpeded by folded Titin I27 domains and operates as a translocation-limited motor, indicating that Hsp104 has evolved a fundamentally different mechanical strategy, one that maintains constant force output regardless of substrate structure. A motor that stalls at each structured element, like ClpB, is poorly suited to processively extract polypeptides from cross-β amyloid and prion fibrils, which are stabilized by extensive hydrogen-bonding networks and pose far greater mechanical resistance than disordered aggregates (*6, 36, 57, 59*). The constant-force, translocation-limited mechanism of Hsp104 may thus represent an evolutionary adaptation that enables sustained processivity through substrates of widely varying mechanical stability (*6, 13, 36, 57, 59*).

We discovered a persistent hyperextended state of Hsp104 hexamers that was not detected in prior work (*28, 33*). This state may have eluded detection because existing cryo-EM reconstructions captured Hsp104 either substrate-free or engaged with soluble casein (*28, 33*), but not while actively remodeling mechanically restrained substrates or aggregates, conditions that may favor or stabilize the hyperextended conformation. Alternatively, the hyperextended state may be transiently sampled under those conditions but at occupancies below the threshold of cryo-EM 3D classification. This newly revealed conformation expands the dynamic range of Hsp104 ring structures. Structural modeling suggests that the hyperextended state can be represented by a composite in which subunits P1–P5 retain the substrate-engaged extended spiral (5VYA, ATPγS-bound) while subunit P6 adopts the more displaced position observed in the ADP-bound open state (5VY8), producing a wider seam opening than captured in either parent structure (Fig. 6).

The hyperextended state raises a key structural question: how do hexamers execute non-sequential conformational transitions, including direct switches between closed and hyperextended states, without disrupting the ring or losing substrate engagement? We propose that ATP hydrolysis at individual protomers provides the energetic driving force: the transition from ATP- to ADP-bound states at a given subunit would weaken inter-protomer contacts locally, enabling the seam to widen beyond the extended state and access the hyperextended conformation (Fig. 6). The threaded polypeptide substrate itself may further stabilize the expanded hexamer by maintaining pore-loop contacts across the ring, effectively acting as a central tether that permits large-scale seam expansion without ring disassembly. Notably, time-resolved cryo-EM of the 26S proteasome AAA+ motor has likewise uncovered non-symmetric conformational changes during substrate processing, with multiple spiral-staircase registers and asymmetric transitions occurring in bursts (*65*). Unlike the heterohexameric proteasomal motor, which comprises six distinct Rpt subunits, Hsp104 achieves comparable conformational plasticity as a homohexamer, indicating that such stochastic dynamics do not require subunit heterogeneity. Whether protomer exchange also contributes to these dynamics under physiological conditions remains an open question. Our observation that translocation proceeds after removal of free Hsp104 from solution (Fig. 1D) demonstrates that protomer exchange is not required, but does not exclude the possibility that subunit exchange in the cellular environment could accelerate recovery from stalled or non-productive states (*6*).

The frequent sampling and prolonged occupancy of extended and hyperextended states across diverse substrates and nucleotide conditions argue for an updated view of AAA+ motor function that explicitly incorporates conformational plasticity. Since each of the six protomer–protomer interfaces can independently adopt any of the three conformations, individual Hsp104 hexamers are likely conformationally heterogeneous at any given moment, with the majority of interfaces typically in extended or hyperextended states and a minority in the closed state. Our findings establish biased stochastic dynamics as a core feature of Hsp104 function rather than an incidental behavior. Sampling conformations probabilistically along preferred pathways allows Hsp104 to preserve substrate engagement and processivity while retaining the versatility needed to bypass otherwise intractable mechanical barriers. That the conformational cycling rate scales proportionally with translocation velocity and disaggregase activity across all conditions (Table S1) further supports tight mechanistic coupling between these dynamics and productive substrate processing. More broadly, these insights expand the mechanistic paradigm of AAA+ ring proteins, illustrating how biased stochastic, energy landscape-driven cycling can support robust and versatile substrate remodeling across the superfamily, and suggest that rigid, strictly deterministic mechanisms would be prone to stalling or failure when confronted with the diverse stabilities and complex geometries of physiological substrates.

These insights lay a mechanistic foundation for engineering disaggregases with enhanced activity or altered specificity. Potentiating mutations (*8, 10–12, 17, 18, 20, 24*) in Hsp104 may act by modulating dwell times, transition biases, or energy barriers between conformations, possibilities that warrant direct testing in future work. The impact of collaborating chaperones such as Hsp70 and Hsp40 on this dynamic cycle also remains an open question, with implications for mechanistic understanding and therapeutic modulation (*10*). The ability to reverse direction, remain engaged with substrates, and dynamically adjust the translocation path in response to barriers defines Hsp104 as a particularly versatile disaggregase (*6*). These features may be especially important for resolving pathogenic protein assemblies in neurodegenerative disease, including aggregates with highly resistant structural cores. Understanding and tuning this mechanistic adaptability could guide the development of next-generation strategies to restore proteostasis and counter neurodegenerative pathology. More broadly, the expanded functional logic of AAA+ motors defined here, in which biased stochastic cycling replaces rigid rotary stepping, may inform the design and engineering of other molecular machines that must navigate complex, heterogeneous substrates.

## Acknowledgments

We thank Zarin Tabassum, Zhuoyi Chen, Raju Roy, Miriam Linsenmeier, and Linamarie Miller for constructive feedback.

## Author contributions

Conceptualization: J.L., D.R.S., A.L.L., Y.E.G., J.S.

Data curation: J.L., J.K.B., A.S., K.A.S., A.L.L., Y.E.G., J.S.

Formal analysis: J.L., J.K.B., A.S., K.A.S., S.P., A.L.L.

Funding acquisition: J.L., S.P., D.R.S., A.L.L., Y.E.G., J.S.

Investigation: J.L., J.K.B., A.S., K.A.S., S.P., D.R.S., A.L.L., Y.E.G., J.S.

Methodology: J.L., J.K.B., A.S., K.A.S., S.P., D.R.S., A.L.L., Y.E.G., J.S.

Project administration: J.L., J.K.B., S.P., A.L.L., Y.E.G., J.S.

Resources: J.L., J.K.B., A.S., K.A.S., S.P., A.L.L., Y.E.G., J.S.

Software: J.L., A.S., K.A.S., S.P., Y.E.G.

Supervision: S.P., A.L.L., Y.E.G., J.S.

Validation: J.L., J.K.B.

Visualization: J.L., J.K.B., A.S., S.P., A.L.L., Y.E.G., J.S.

Writing – original draft: J.L., J.K.B., A.S., S.P., A.L.L., Y.E.G., J.S.

Writing – review & editing: J.L., J.K.B., A.S., K.A.S., S.P., D.R.S., A.L.L., Y.E.G., J.S.

## Data and materials availability

Plasmids generated in this study will be made readily available to the scientific community. All requests will be honored in a timely manner. All data used for this study are available in the manuscript, the supplementary material, or deposited at indicated data repositories. The paper does not report original software, but custom Python scripts and ImageJ analysis code are available on Zenodo (doi: 10.5281/zenodo.18295617).

## Materials and Methods

### Materials

#### Site-directed mutagenesis

Mutations in Hsp104, MFSV_S736C (C209V:C400A:C643M:C718F:C721S:S736C:C876V),

SASV_S736C (C209V:C400A:C643S:C718A:C721S:S736C:C876V), MASV_E793C

(C209V:C400A:C643M:C718A:C721S:E793C:C876V), MFSV_E793C

(C209V:C400A:C643M:C718F:C721S:E793C:C876V), MFSA_E793C

(C209V:C400A:C643M:C718F:C721S:E793C:C876A), SFSV_E793C

(C209V:C400A:C643S:C718F:C721S:E793C:C876V), MASV_E793C:N728A and MASV_E793C:E360R were introduced using the QuikChange site-directed mutagenesis kit (Agilent Technologies) and confirmed by Sanger DNA sequencing.

### Generation of vectors for protein purification

#### β-casein-AviTag vector construction

A synthetic gene block encoding β-casein with a C-terminal cysteine was obtained from Integrated DNA Technologies (IDT). The pMal-T-Avi-HisBirA vector (Addgene: 102962) was linearized by polymerase chain reaction (PCR) with specific primers. The β-casein gene block was inserted into the linearized vector using Gibson assembly (New England Biolabs) to create a construct encoding an open reading frame (ORF): MBP-Factor Xa-β-casein-AviTag-HisTag. The final construct was verified by sequencing.

#### Sup35-NM-AviTag Vector Construction

The pAED4-Sup35-NM-A230C vector (Addgene: 15598) was linearized by PCR with specific primers. An AviTag was inserted via Gibson assembly to generate an ORF encoding Sup35-NM-A230C-AviTag-HisTag. The construct sequence was verified. The pH6-MBP-TEV-BirA vector (Addgene: 179694) was used for the purification of MBP-TEV-BirA for biotinylation.

### Protein purification

#### Hsp104 Purification and Labeling

Hsp104^WT^, Hsp104^N728A^, and single-cysteine variants Hsp104: SASV_S736C, MASV_E793C, and MASV_E793C:N728A (see fig. S2) were expressed and purified as described previously (*10*). Proteins were labeled with Cy3 or Cy5 maleimide dyes (Sigma-Aldrich). After labeling, free dye was removed via 6-hour dialysis, followed by purification using a HiTrap Desalting column (GE Healthcare). Labeling efficiency was quantified using a UV-visible spectrometer. The extinction coefficient 314000 M^-1^ cm^-1^ (monomeric) was used to determine Hsp104 concentration. For conformational change experiments, labeled and unlabeled Hsp104 were mixed in ratios of 1:1:4 or 2:2:2 (Cy3-labeled: Cy5-labeled: unlabeled).

#### β-casein-AviTag-His Expression and Protein Purification

The pMal-T-Avi-His vector encoding MBP-Factor Xa-β-casein-AviTag-HisTag was transformed into *E. coli* BL21 (DE3)-RIL cells. Cultures were grown in 2xYT medium supplemented with 100 µg/ml ampicillin and 50 µg/ml chloramphenicol at 37°C until OD_600nm_ reached ∼0.6–0.8. Protein expression was induced with 1 mM IPTG supplemented with 0.05 mM biotin for *in-vivo* biotinylation. Cultures were incubated overnight at 16°C. Cells were harvested by centrifugation (4,658×g, 4°C, 25 min), resuspended in lysis buffer (40 mM HEPES-KOH pH 7.4, 20 mM imidazole, 20% glycerol, 2 mM β-mercaptoethanol, 0.05 mM biotin, EDTA-free-Complete-protease inhibitors), lysed by sonication, and centrifuged (30,996×g, 4°C, 20 min).

The supernatant was incubated with Ni-NTA beads with shaking for 3 hours, washed with lysis buffer, and eluted with 40 mM HEPES-KOH pH 7.4, 500 mM imidazole, and 2 mM β-mercaptoethanol. Eluted protein was subjected to gradient purification using a HisTrap FF column with a 20-column volume gradient of imidazole (20–500 mM). SDS-PAGE analysis confirmed >99% purity. Biotinylation efficiency was validated using the Pierce Biotin Quantification Kit. MBP-Factor Xa-β-casein-AviTag-HisTag was cleaved with Factor Xa (1:1000 ratio) overnight. Cleaved β-casein-AviTag-His was resuspended in HKM150 (50 mM HEPES pH 7.4, 150mM potassium acetate, 20 mM Mg(OAc)₂, 2 mM β-mercaptoethanol) with 6 M urea, labeled with Cy5 maleimide, and purified using sequential Micro-Bio-Spin P-6 columns (Bio-Rad). Labeling efficiency was confirmed by UV-visible spectrometry.

### RepA-Titin_n_ purification

RepA-Titin_1_, RepA-Titin_2_, and RepA-Titin_3_ were purified and labeled with AF555 as described (*51, 52*).

#### Sup35-NM-A230C-AviTag-His Protein Expression and Purification

The pAED4-Sup35-NM-A230C-AviTag-His and pH6-MBP-TEV-BirA vectors were independently transformed into *E. coli* BL21 (DE3)-RIL cells. Cultures were grown in 2×YT medium containing 100 µg/ml ampicillin and 50 µg/ml chloramphenicol at 37°C until OD_600nm_ reached ∼0.6–0.8. Protein expression was induced with 1 mM IPTG supplemented with 0.05 mM biotin. Cultures were incubated overnight at 16°C. Cells were harvested, lysed as described for β-casein, and the supernatant was incubated with Ni-NTA beads for 3 hours. Proteins were eluted with 40 mM HEPES-KOH pH 7.4, 500 mM imidazole, 2 mM β-mercaptoethanol, and 0.05 mM biotin.

Eluted protein was further biotinylated (with 5 mM biotin) overnight at 4°C. Aggregated Sup35-NM-A230C-AviTag-His was separated from soluble His-MBP-TEV-BirA by centrifugation and resuspended in 40 mM HEPES pH 7.4, 8 M urea. Purity was assessed by SDS-PAGE, confirming >95% purity. The biotinylated protein was labeled with Cy5 maleimide in 8 M urea and purified using sequential Micro-Bio-Spin P-6 columns to check its specificity to streptavidin binding. Labeling efficiency was verified by UV-visible spectrometry.

#### Sup35-NM-A230C-AviTag-His fibril preparation

247 µM Sup35-NM-A230C-AviTag-His protein (in 40 mM HEPES, pH 7.4, 8 M urea) was diluted into HKM150 to a final concentration of 1.2 µM and incubated overnight at 4 °C with agitation at 25 rpm. Fibril formation was then assessed under bright-field microscopy at 100× magnification prior to the performance of single-molecule experiments.

## Methods

### Generative REgularized ModeLs of proteINs (GREMLIN) Coevolution Analysis

GREMLIN coevolution analysis was performed using OPENSEQ.org web server (*67*). The primary sequence of *Saccharomyces cerevisiae* Hsp104 was used as input. Multiple sequence alignments (MSA) were generated using HHblits with an E-value cutoff of 10^-10^ and four interaction iterations. Filtering was applied to remove sequences covering less than 75% of the query, and positions with more than 75% gaps were excluded.

### Thermotolerance assay

pAG416GAL plasmids containing vector control, Hsp104^WT^ and the single-cysteine variants, MFSV_S736C, SASV_S736C, MASV_E793C, MFSV_E793C, MFSA_E793C, and SFSV_E793C, under the GAL promoter were transformed into W303a*Δhsp104* yeast. Cultures were grown overnight in raffinose dropout media at 30°C. Cells were normalized to OD_600nm_ = 0.3, shifted to galactose dropout media for 5 hours to induce expression, and normalized to OD_600nm_ = 0.6. Heat shock was performed at 50°C for 10, 20, and 30 minutes in a thermomixer, followed by 2 minutes on ice. Five-fold serial dilutions were plated on glucose (no induction) and galactose (induction) agar plates and incubated at 30°C for 3 days before imaging.

### Luciferase Disaggregation and Reactivation Assays

Luciferase aggregates were prepared by incubating 6 mg/mL recombinant firefly luciferase (Sigma-Aldrich) in HKM150-LF buffer (25 mM HEPES-KOH pH 7.4, 150 mM potassium acetate, 10 mM magnesium acetate, 10 mM DTT) supplemented with 6 M urea at 30°C for 30 minutes. The denatured luciferase was diluted 100-fold on ice into HKM150-LF buffer (without urea), snap-frozen, and stored at −80°C.

Disaggregation assays were performed as described previously (*8, 48*). Briefly, 1 µM Hsp104, with or without 0.167 µM Hsc70 and 0.167 µM Hdj2 (*68*), was incubated with 100 nM luciferase aggregates (monomeric concentration) in the presence of an ATP regeneration system (ARS: 20 mM creatine phosphate, 5 mM ATP, 20 μg/mL creatine kinase) in HKM150-L buffer. Samples were incubated at 25°C for 90 minutes, mixed with luciferase assay reagent (Promega), and luminescence was measured using a Safire Tecan luminometer.

### Single-Molecule TIRF Experimental Setup

#### Slide Preparation and Illumination Buffer

Slides were prepared following an adapted protocol for protein passivation (*69*) with some modifications (*70, 71*). Channels were washed with 20 µL 1% Tween in HKM150 buffer for 5 minutes and rinsed with HKM150. To verify passivation quality, one channel was loaded with 0.5 mg/mL streptavidin, incubated for 2 min, and washed with 20 µL HKM150; 10 nM Cy5-casein-biotin was then loaded into this channel and into a parallel channel lacking streptavidin. A specific-to-nonspecific binding ratio far exceeding 100:1 was confirmed prior to experiments. To minimize photobleaching, an enzymatic oxygen scavenging system was included: 0.3% (w/v) glucose, 300 μg/mL glucose oxidase (Sigma-Aldrich), 120 μg/mL catalase (Roche), and 1.5 mM Trolox (Sigma-Aldrich) (*72*). This enzymatic deoxygenation system will be referred to as the IB (illumination buffer) system in the text.

#### Measurement of Hsp104 Translocation

Biotinylated Cy5-labeled β-casein substrates were immobilized via biotin-streptavidin interactions. Cy3-labeled Hsp104 and unlabeled Hsp104 were mixed to final concentrations of 10–20 nM (labeled) and as indicated in the text (unlabeled). Imaging was performed at room temperature (24.6–25.8°C) on a custom-built objective-type TIRF microscope (Nikon Eclipse Ti-E) (*72*), with 532 nm and 640 nm laser excitation for Cy3 and Cy5, respectively. Experiments were conducted in HKM150 or HKM25 buffer (50 mM HEPES pH 7.4, 150 or 25 mM potassium acetate, 20 mM Mg(OAc)_2_, 2 mM β-mercaptoethanol). Higher HEPES and Mg(OAc)_2_ concentrations (compared to the HKM buffer used in the Luciferase disaggregation and reactivation assays) were used to limit potential pH shifts caused by illumination buffer (IB) components during single-molecule experiments. Image stacks were recorded at 10 or 25 frames/s using custom LabView scripts, which also triggered laser shutters and pumps and are available under a CC-BY-ND 4.0 International license.

#### Measurement of Hsp104 Conformational Changes

A 3 µM (monomeric) mix of Hsp104^N728A^ or Hsp104 (Cy3-Hsp104:Cy5-Hsp104:unlabeled Hsp104 = 1:1:4 or 2:2:2) was incubated with 5 mM ATP (with ARS) or ATP:ATPγS (1:1, 2.5 mM each) for 10 min, then diluted to 60–600 nM Hsp104 (monomeric concentration) plus 5 mM ATP (with ARS) or ATP:ATPγS and IB components immediately before loading into the observation channel. A 20 µL aliquot of this sample was loaded into the control channel, incubated for 2 min, and rinsed with 20 µL HKM150 or HKM25 containing 5 mM of the corresponding nucleotide plus IB, before smTIRF imaging.

To measure Hsp104 conformational dynamics, a separate channel on the same slide was loaded with 20 µL casein (20–600 nM; 1:1 molar ratio relative to the Hsp104 hexamer) or Sup35-NM fibrils (20–120 nM monomeric concentration, diluted immediately before loading from a 1.2 µM pre-assembled stock), incubated for 5 min, and rinsed. The Hsp104-nucleotide mix plus IB (20 µL) was then loaded, and smTIRF recording began immediately. Cy3 and Cy5 fluorescence were collected by alternating laser excitation (ALEX), switching between 532 nm and 640 nm excitation every other frame, with a 100 ms integration time per color (200 ms total frame rate). FRET traces were analyzed using custom software (71), implemented in ImageJ or Python scripts available on Zenodo (<u>ImageJ_Python_scripts).</u>

The distance between the donor (Cy3) and the acceptor (Cy5), *r,* can be determined from the FRET efficiency by the Förster equation (Equation 1):

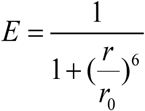

Where the Förster distance, *r_0_,* for the Cy3-Cy5 FRET pair is 5.6 nm.

FRET efficiency determined from TIRF microscope experiments was calculated using Equation 2:

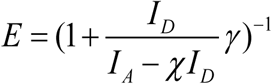

where *I_D_* and *I_A_* are the background corrected fluorescence intensities of the donor and acceptor, χ is the crosstalk of the donor emission into the acceptor recording channel, and γ accounts for the ratios of quantum yield and detection efficiency between the donor and the acceptor channels. Those correction factors were determined for each experiment as previously described (*70, 72, 73*).

### Single-turnover stopped flow determination of Hsp104 unfolding and translocation

Hsp104 and RepA-Titin_n_ were dialyzed into HK150 buffer (25 mM HEPES, 150 mM KCl, 10 mM MgCl_2_, 2 mM dithiothreitol, and 10% glycerol (v/v) with pH 7.5 at 25°C) using 50 kDa and 10 kDa molecular weight cutoff dialysis tubing, respectively. The experiments were performed in an SX20 Applied Photophysics stopped-flow fluorometer (Leatherhead, U.K.) with a sequential mixing set-up at 25 °C. The mixing schemes and detailed reagents are illustrated in Fig. 3A, B. Previous work (*51, 52*) provides more details on the experimental strategy.

#### Sample Preparation

Hsp104 at a concentration of 4 µM was loaded into Syringe 1, while Syringe 2 contained a mixture of ATPγS as indicated in the text and 200 nM RepA-Titin_n_ substrates with repeat lengths of *n* = 1, 2, or 3. The contents of Syringes 1 and 2 were rapidly mixed in a reaction chamber and incubated in an aging loop for a user-defined delay time (Δt₁). During Δt₁, Hsp104 formed stable hexamers in the presence of ATPγS and bound to the RepA sequence of the RepA-Titin_n_ substrate.

#### Reaction Initiation and Monitoring

After the Δt₁ incubation period, the aged mixture was rapidly mixed with α-casein and ATP at concentrations indicated in the text to initiate the translocation and unfolding reactions. Protein-induced fluorescence enhancement of AF555 was monitored over time (Δt₂) using a fluorescence detection system. The α-casein acted as a trap for released Hsp104 to ensure observations corresponded to single-turnover translocation of Hsp104 on RepA-Titin_n_.

#### Data Collection and Kinetic Analysis

Time-course fluorescence data were collected at pre-incubation times (Δt₁) of 120, 360, and 600 seconds for each RepA-Titin_n_ repeat length. The translocation and unfolding rates were determined by plotting the total length of the RepA-Titin_n_ substrate against the time to reach peak fluorescence for each Δt₁. Linear regression analysis of these data provided the rate of translocation and unfolding at each Δt₁. To estimate the translocation/unfolding rate at Δt₁ = 0, rates obtained at varying Δt₁ values were plotted as a function of Δt₁, and the intercept of the linear fit was calculated.

#### ATPγS-Dependent Rate Analysis

To assess ATPγS-dependent dynamics, the length of amino acids excluded from translocation/unfolding during the ATPγS phase was plotted as a function of Δt₁. The slope of this plot represents the ATPγS-driven unfolding rate, while the intercept corresponds to the length of the amino acid segment not engaged in translocation. These segments likely included amino acids located on the exterior of the hexamer without interacting with the NBDs. The decrease in translocation and unfolding rates with increasing Δt₁ suggested hydrolysis of ATPγS to ADP during the pre-incubation phase, leading to higher ADP concentrations that inhibited Hsp104 activity. These observations provided insights into the nucleotide-dependent dynamics of Hsp104-mediated substrate translocation and unfolding.

### Bayesian analysis to determine the number of Hsp104 conformational states

The raw donor and acceptor fluorescence intensities I_D_ and I_A_ were converted to FRET efficiencies, *E*, using Equation 2. Time points with fluorescence intensity values less than zero due to noise were omitted from the calculation of *E* but flagged so as to preserve the correct time stamps for all the data. FRET efficiencies were then analyzed with a Bayesian mixture model in order to determine the most likely number of FRET states. The analysis was adapted from a previous method used to identify the number of different motional states of side chains from NMR-derived methyl order parameters (*54*). FRET efficiencies were assumed to report on an unknown number of conformational states, *M*, each with an intrinsic value of FRET efficiency, *E*_j_, plus Gaussian-distributed noise with root mean square standard deviation σ. A single time series of FRET measurements consists of a sampling from this mixture of states, where each state is occupied for some average fraction *f*_j_ of the total experimental time, where

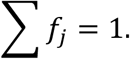

The likelihood of a set of FRET efficiency measurements {*e_k_*}, *k* = 1, N in a Gaussian noise model is given by (Equation 3)

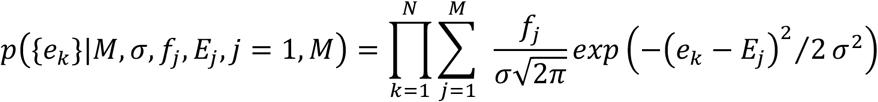

Using the global average values of intrinsic efficiency for states 1, 2, and 3, the values midway between 1 and 2, and 2 and 3 are 0.40 and 0.65 for each experiment, respectively. In the Gaussian noise model, a state with a FRET efficiency equal to the midpoint value has an equal probability of coming from either states 1 and 2 or 2 and 3, respectively. Thus, one can assign each FRET efficiency data point to a state as follows. State 1: e*_k_* ≤0.40, state 2: 0.40 < e*_k_* < 0.65, state 3: e*_k_* ≥ 0.65, producing a time series of states for each experiment. These time series were then analyzed to obtain mean residence times and average state occupancies. These are summarized in Table S3.

### Determination of Hsp104 conformational states and their transition kinetics

Time trajectories with anticorrelated donor and sensitized emission were manually selected from the traces identified with FRET pairs (*53*). Trajectories with more than 1 anticorrelated signal change that was not due to photo bleaching were selected. Individual smFRET traces collected within each experimental replicate were concatenated into a single composite trace prior to hidden Markov model (HMM) analysis. This approach addresses a known limitation of three-state HMM analysis applied to short individual traces: label degeneracy, in which insufficient transitions prevent reliable estimation of all three FRET state means. Visual inspection of per-molecule Viterbi state assignments confirmed this behavior, the same conformational state (e.g., FRET ∼ 0.4) was assigned different indices across traces, or unvisited states were placed at physically implausible FRET efficiencies driven by the prior rather than the data. This degeneracy arises from three compounding factors:

- Insufficient transition sampling: short traces may visit only two of the three states, leaving the third FRET mean unconstrained and prior-dominated.
- Multimodal ELBO (Evidence Lower BOund) landscape: with few data points, variational Bayes optimization converges to different local optima (label permutations) across molecules, yielding inconsistent state indices (*53, 74*).
- Prior domination: sparse observations allow the hyperprior on state means to dominate the likelihood, inflating variability in inferred FRET state positions.

By concatenating the recordings from all molecules from a single replicate into one trace, the total number of state transitions increased sufficiently to sharply constrain all three FRET state means within a single optimization, yielding consistent and reproducible state assignments across molecules within each replicate.

To prevent the HMM from inferring spurious transitions at molecule boundaries, an artificial break-marker frame with a FRET value greater than 2 was inserted between consecutive traces. Because the emission probability of a value exceeding 2 is effectively zero under any physically meaningful Gaussian state model (FRET means 0.1 - 0.9), the Viterbi decoder assigns break-marker frames to an undefined state and does not update the transition matrix across boundaries. Break-marker frames were excluded from all downstream dwell and occupancy analyses. This approach is mathematically equivalent to ebFRET’s native multi-trace input mode for state mean estimation, while providing a continuous input format compatible with the software interface HMMs were fit to the concatenated data using ebFRET with model orders K = 2–6 states. A three-state model was selected on the basis of per-trace Bayesian mixture model comparison (Table S3), in which four- and five-state solutions were in every trajectory several orders of magnitude less likely than the three-state description, and corroborated by variational lower bound comparison across model orders (Tables S4, S5), which showed the largest gain in approximate model evidence at the 2→3 state transition with progressively diminishing returns thereafter. State trajectories were decoded by the Viterbi algorithm, and dwell events were exported for downstream analysis. Restarts and precision of 10^-2^ were set for the initial iteration and 10^-6^ for the final reiteration. The global analysis constrained the FRET efficiency states, and the Viterbi Mean of the FRET efficiency to be the same for all replicates. 5-6 replicates were performed for each Hsp104-substrate condition. A MATLAB program was used to generate the transition density map and dwell time distribution (*53*). GraphPad Prism was used to plot the histogram distributions and cumulative dwell time distributions which were fit to single-exponential curves using GraphPad Prism with Equation 4:

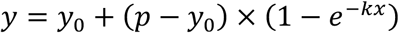

Here *p* is the plateau of the distribution and *k* is the estimated rate constant. The half-time is calculated: *T*_1/2_ = ln (2)/*k*.

For the subpopulation analyses shown in Figs. S5, S7, and S8, trajectories were grouped by the number of conformational states they sampled and fitted with ebFRET with the number of states held fixed at the category value, to resolve the mean FRET efficiency and dwell-time mode of each state within each subpopulation. Category assignment was initialized from the per-trace Bayesian mixture model (Table S3) and refined against the fixed-state ebFRET fit: trajectories for which the fit under the initially assigned state number did not adequately describe the data, for example, a fitted state that did not coincide with an occupied FRET level, or a resolved transition left unmodeled, were reassigned to the state number whose ebFRET fit was consistent with the trajectory, improving the agreement between the assigned model and the observed FRET behavior. This refinement is therefore semi-independent of Table S3, seeded by the per-trace classification and adjusted by trajectory-level fit quality. Net reassignment was directional in every condition, moving trajectories with a resolvable but low-occupancy third FRET level from the 2-state to the 3-state category (WT Hsp104 on Sup35-NM: 7 of 169 trajectories, ∼4%; Hsp104-N728A on β-casein: 7 of 105 trajectories, ∼7%; WT Hsp104 on β-casein: 19 of 149 trajectories, ∼13%); the dominant three-state category was preserved under both classifications. Among hexamers that sampled only two states, the hyperextended-extended subpopulation (State 1→2) was the more common across conditions.

### Model order selection and Bayes factor analysis

To determine how many discrete conformational states are justified by the data, we used the variational lower bound reported by ebFRET as an approximate log marginal likelihood (log-evidence) for each candidate model order. For each condition (WT–β-casein, N728A-β-casein, and WT-Sup35-NM fibrils), ebFRET was run with model orders K = 2,…,6 states and the resulting summary “2-6 states” CSV files were exported. In these files, the Lower_Bound section reports, for each K, the mean lower bound per series, which provides a goodness-of-fit measure averaged over all trajectories for that condition.

Interpreting the mean lower bound ℒ*_K_* as an approximation to log *p*(data ∣ *K*), differences Δℒ = ℒ*_K’_* − ℒ*_K_* correspond to approximate log Bayes factors comparing model orders *K*^’^ and *K*:

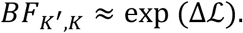

We therefore used the differences in mean lower bound between K = 2 and higher-order models (K = 3-6) to assess the strength of evidence for additional conformational states, adopting the conventional Jeffreys/Kass–Raftery scale in which Δℒ ≳ 5 (i.e., *BF* ≳ 150) is considered “decisive” support (*75*). Because the lower bounds are reported on a per-series basis, they already reflect averaging across traces, and the reported values represent the typical evidence gain per hexamer on moving from 2-state to higher-state models.

### Quantification of conformational state occupancy from smFRET dwell-time data

State occupancies were quantified from ebFRET-derived (*53*) dwell-time transition files, in which each row records a single dwell event comprising the starting FRET state, the subsequent transition target, and the dwell duration in the starting state measured in camera frames. For each condition, FRET values were assigned to one of three discrete states using midpoint thresholds between the ebFRET-fitted state FRET efficiencies.

Fractional state occupancy for each replicate *i* was defined as the percentage of analyzed frames assigned to a given state *s*. Specifically, frames_i_(*s*) denotes the total number of frames corresponding to dwell events originating from state *s*, and totalframes_i_ is the total number of frames analyzed in replicate *i*. Occupancy was therefore calculated as:

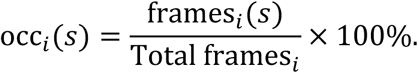

Because replicate sizes varied substantially within the N728A condition (507–22,636 total frames per replicate), state occupancies *x_i_*_i_ (where *x_i_*_i_ = occ*i*(*s*)) were aggregated using a frame-weighted mean. Each replicate was assigned a weight *w_i_*. proportional to its total analyzed frames frames. (i.e., frames*_i_* ≡ Total frames*_i_*), normalized by the average frame count across replicates:

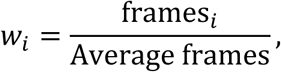

such that the sum of weights satisfies ∑_i_*w*_i_ = *n*, where *n* is the number of replicates. The weighted mean occupancy *x^-^_w_* was then computed as:

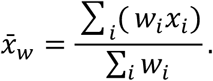

This weighted mean is equivalent to the pooled occupancy obtained by combining all frames across replicates.

To estimate uncertainty, we calculated a Bessel-corrected weighted variance, where (*x_i_*−*x^-^_w_*) represents the deviation of each replicate’s occupancy from the weighted mean:

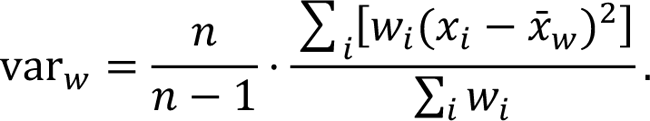

The weighted standard error of the mean (SEM) was then defined as:

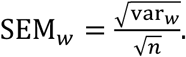

### Bootstrap estimation of 95% confidence intervals for transition probabilities

#### Construction of discrete state sequences and transition matrices

For each of the three experimental conditions, N728A-β-Casein in ATP, WT-β-Casein in ATP-ATPγS, and WT-Sup35-NM in ATP, ebFRET analysis exported transition probabilities were subjected to Bootstrap estimation of the confidence interval (Table S7). In the input data, each row represented a dwell segment and its subsequent transition, with columns specifying a trace identifier, the FRET value before the transition, the FRET value after the transition, and the dwell duration in frames.

To obtain a consistent three-state representation per condition, continuous FRET values were mapped to the nearest of three condition-specific centers determined by ebFRET:

- WT-β-Casein in ATP-ATPγS: 0.27 (state 1, low FRET), 0.50 (state 2, mid FRET), 0.69 (state 3, high FRET).
- N728A-â-Casein in ATP: 0.23 (state 1), 0.45 (state 2), 0.74 (state 3).
- WT-Sup35-NM in ATP: 0.18 (state 1), 0.43 (state 2), 0.65 (state 3).

For each dwell, we assigned a discrete “from-state” and “to-state” (1, 2, or 3) by mapping the corresponding FRET values to the nearest center. Self-transitions (i→i) were excluded, as they reflect persistence in a state rather than a transition, and all remaining non-self transitions between consecutive dwells were collected as a list of individual events.

For each condition, we constructed a 3×3 transition count matrix *N*, where *N_ij_* is the number of observed transitions from state i to state j (i ≠ j). Row sums *N_i_ = Σ_j≠i_ N_ij_* gave the total number of leaving events from origin state i. The empirical transition probability matrix *P* was defined by row normalization:

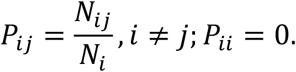

These row-normalized matrices provided the point estimates of the per-edge transition probabilities for each condition.

#### Nonparametric bootstrap of individual transition events

To quantify uncertainty in the estimated transition probabilities, we treated the list of individual non-self transitions as the basic sampling unit and performed a nonparametric bootstrap. For a given condition, let (*i_k_^’^, j_k_*) denote the origin and destination state of the k-th non-self transition in the dataset, for k = 1,…,N.

We generated B bootstrap replicates (B = 5000) as follows:

1. **Resampling:** Draw N transition indices with replacement from {1,…,N} to form a bootstrap sample of transitions *{(i^(b)^_k_,j^(b)^_k_)}^N^_k=1_*.
2. **Bootstrap count matrix:** From this resampled set, construct a 3×3 bootstrap count matrix *N*^(*b*)^by incrementing *N^(b)^_ij_* for each occurrence of i→j (i ≠ j).
3. **Bootstrap transition matrix:** Row-normalize *N*^(*b*)^ to obtain a bootstrap transition probability matrix *P*^(*b*)^, with

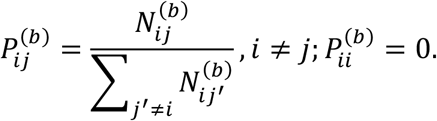

This procedure yields a set of B bootstrap estimates *P^(b)^_ij_* for each directed non-self transition i→j and each condition.

#### Confidence interval estimation

For each condition and for each directed non-self transition i→j, we formed the empirical bootstrap distribution *{P^(b)^_ij_}^B^_b=1_*. The 95% confidence interval (CI) for the transition probability *P_ij_* was obtained using the percentile method: the lower and upper bounds were defined as the 2.5th and 97.5th percentiles of the bootstrap distribution, respectively:

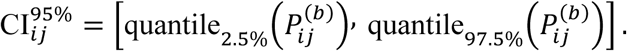

The point estimate reported in the main text for each transition was the row-normalized probability from the original (non-bootstrapped) data, and the associated 95% CI was taken from these bootstrap percentiles.

### Model-independent simulation of Hsp104 conformational change

To simulate a realistic smFRET signal for comparison with the experimental results, we first used the classic Gillespie algorithm (*76*) to generate a state trajectory {*s*_1_, *s*_2_, *s*_3_, … } with dwell times {Δ*s*_1_, Δ*s*_2’_, Δ*s*_3_, … }, based on a given kinetic scheme represented by a transition rate matrix Λ. Next, the trajectory was divided into equally-sized time bins based on the experimental frame rate, where we assumed average dwell times in each state were longer than the frame interval. Then, based on the typical total photon count per bin and FRET efficiencies determined by the state trajectory, we calculated sample donor and acceptor emissions. Finally, EMCCD camera noise was added to the donor and acceptor counts by applying realistic Gamma-distributed electron-multiplication with a gain of 40, readout sensitivity of approximately 5 electrons per digital count, as well as normally-distributed readout noise of approximately 2 electrons with a base offset of 150 digital counts, all integrated over a 5×5 pixel grid surrounding the donor/acceptor.

More formally, the transition rate matrix for a six-state model with states labeled as {σ_1_, σ_2_, …σ_3_} is given by

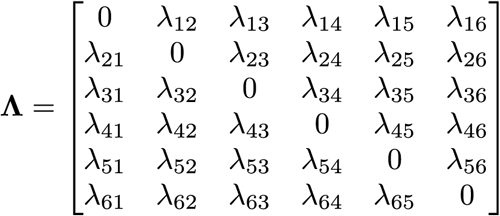

where *λ_ij_* is the transition rate for switching from states *σ_i_* to *σ_j_*. Next, to generate the state trajectory, we first initialized the trajectory by a randomly chosen state *s*_1_. The dwell time for the initial state was then sampled as

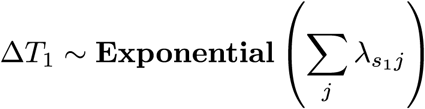

and the next state was sampled from a Categorical distribution, which stochastically selects one of the six states based on a probability vector (that sums to 1) determined by transition rates, as below

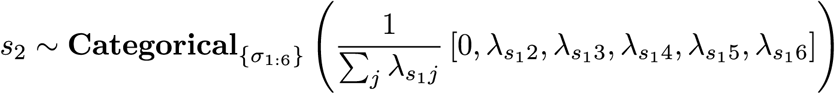

where the self-transition probability is always set to 0 (as shown above for demonstration when *s*_1_ = σ_2_). This process was repeated until the temporal length of the state trajectory {*s*_1_, *s*_2_, *s*_3_ …} becomes longer than the typical duration of the experiment.

Next, the state trajectory was discretized into N time bins marked by a discrete set of time points {*t*_0_,*t*_1_,…,*t_n_*,…,*t_N_*} where the n-th bin lies between *t_n_*_–1_ and *t_n_*. The FRET efficiencies {*E*_0_,…,*E_n_*,…,*E_N_*} were then assigned to each time bin based on the state trajectory.

Now, assuming an average donor laser excitation rate of *λ^ex^* photons per bin, we sampled the Poisson-distributed number of donor photon absorptions as

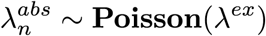

which result in FRET efficiency dependent donor and acceptor photon emissions sampled as

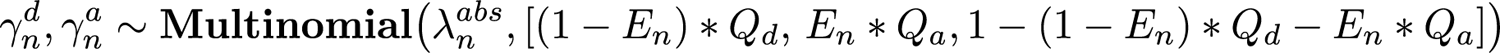

where *Q_d_*, *Q_a_* are donor and acceptor quantum yields, respectively, and multinomial distribution was chosen to enforce the number of total events being equal to total absorptions.

Next, the emitted photons entering the detector generate photoelectrons which are multiplied at the EM gain stage of the EMCCD camera to produce Gamma-distributed multiplied electrons in the donor and acceptor channels as (*77*).

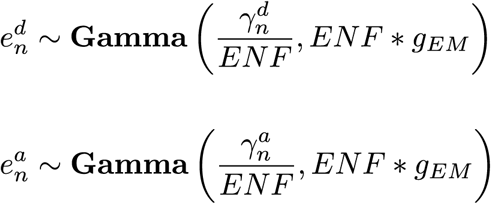

where *g_EM_* and ENF≈2.0 are EM gain and excess noise factor(*77*), respectively.

Finally, normally-distributed readout noise was applied to these multiplied electrons to give donor and acceptor channel digital counts *d_n_*, a_n_ as

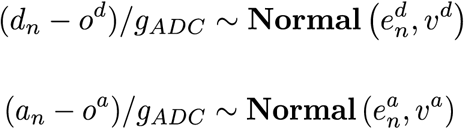

where *o^d^*, *o^a^* and *υ^d^*, *υ^a^* are camera readout offsets and variances for the donor and acceptor channels, respectively. The analog-to-digital readout sensitivity *g_ADC_* was selected on the camera such that the maximum brightness of the signal did not saturate pixels resulting in erroneous counts.

## Supplementary Text

### Engineering a singly labeled Hsp104 variant with WT activity

To innovate a singly labeled Hsp104 variant with activity comparable to WT Hsp104, cysteine–maleimide conjugation chemistry was used. First, the six native cysteines in *Saccharomyces cerevisiae* Hsp104 were replaced and a single cysteine was introduced at a surface-exposed site suitable for FRET measurements without compromising activity. GREMLIN conservation analysis across 7,859 homologs indicated that none of the six cysteines of *S. cerevisiae* Hsp104 are highly conserved (fig. S2A). Guided by this analysis, Hsp104 variants were engineered by substituting each cysteine with the most frequently observed residue at that position or with serine. One cysteine-light background, containing substitutions 209C→V, 400C→A, 643C→S, 718C→A, 721C→S, and 876C→V, is referred to as VA<u>SASV</u> (abbreviated to SASV, as V and A were fixed at positions 209 and 400, respectively, in all designs described here). Two solvent-exposed positions, S736 and E793, were strategically selected for site-specific cysteine insertion and fluorophore labeling to enable tracking of Hsp104 and the spiral seam (fig. S2A–C).

The functional activity of cysteine-substituted Hsp104 variants was assessed in *S. cerevisiae*, where Hsp104 is required for survival under heat shock (*78*). In thermotolerance assays, W303a*Δhsp104* strains expressing single-cysteine Hsp104 variants displayed viability comparable to WT Hsp104 after exposure to 50°C for 0, 10, 20, or 30 minutes (fig. S2B). Specifically, the SASV-S736C and MASV-E793C (209C→V, 400C→A, 643C→<u>M</u>, 718C→<u>A</u>, 721C→<u>S</u>, and 876C→<u>V</u>) variants conferred WT-level thermotolerance (fig. S2B). These variants were purified, site-specifically labeled with Cy3 or Cy5, and tested for disaggregation and reactivation of luciferase trapped in chemically-denatured aggregates in the presence and absence of Hsc70 (an Hsp70) and Hdj2 (an Hsp40; fig. S2D and Methods). The MASV-E793C-Cy3:Cy5 variant retained WT-like disaggregation activity, whereas SASV-S736C-Cy3:Cy5 showed ∼40% reduced activity (fig. S2D). Thus, MASV-E793C was selected as the preferred background for subsequent experiments.

To enable direct comparison with Hsp104^N728A^, the N728A mutation was introduced into the MASV-E793C background. The Hsp104^N728A^ variant binds ATP at NBD2 but cannot hydrolyze it, leading to strong substrate engagement (*28, 46, 48*). The resulting MASV-N728A-E793C construct reactivated luciferase aggregates similarly to Hsp104^N728A^ in both the presence and absence of Hsp70 and Hsp40 (fig. S2D). Notably, MASV-N728A-E793C matched Hsp104^N728A^ in luciferase reactivation, supporting the functional equivalence of MASV-E793C to WT Hsp104. In parallel, the potentiating E360R mutation, which enables mitigation of aggregation and toxicity of neurodegenerative disease-associated proteins (*12*), was introduced into the MASV-E793C background. The MASV-E360R-E793C variant effectively reduced the toxicity of α-synuclein, FUS, and TDP-43 in yeast, indicating that MASV-E793C preserves full functionality (fig. S2E). Together, these results establish MASV-E793C as a robust, functional analog of Hsp104 suitable for single-molecule TIRF microscopy studies of translocation, substrate engagement, and conformational dynamics.

### Hsp104^N728A^ exhibits interaction-site preferences on casein

To measure Hsp104 translocation dynamics, three loading strategies were tested for assembling biotinylated β-casein and Hsp104^N728A^ on slide surfaces (fig. S3A-C). These approaches aimed (1) to stabilize single β-casein–Hsp104^N728A^ complexes, as β-casein can form multimers at nanomolar concentrations in HKM150 (producing large Cy5-labeled clusters), and (2) to assess whether Hsp104^N728A^ shows preferred binding positions along the β-casein sequence under different conditions.

In the first approach, Hsp104^N728A^ and β-casein were preincubated at equimolar ratios for 15–30 minutes at varying concentrations in HKM150 containing 5 mM ATP and an ATP-regenerating system (fig. S3A). This strategy minimized loading of multiple Hsp104 hexamers on a single β-casein molecule and dispersed β-casein–Cy5 clusters, yielding stable single Hsp104–Cy3:β-casein–Cy5 complexes. Approximately 46% of Hsp104^N728A^ hexamers bound at positions corresponding to a FRET efficiency of ∼0.2 relative to the β-casein C-terminus, indicating a preferred interaction site ∼7.1 nm from the C-terminus at equilibrium.

In the second and third approaches, β-casein was first denatured in 8 M urea, diluted to 4 M urea, and then immobilized on the slide before adding Hsp104^N728A^. Measurements were initiated either after preincubating Hsp104^N728A^ with ATP to promote hexamer assembly (fig. S3B) or by adding ATP immediately before recording (fig. S3C). Preincubation with ATP led to ∼41.6% of engagement events initiating at zero FRET efficiency, consistent with a preferred starting position far from the C-terminal region (fig. S3B). By contrast, when ATP was added just prior to data acquisition (fig. S3C), Hsp104^N728A^ showed a broader distribution of initial binding positions, suggesting more frequent side loading onto β-casein (*79*) rather than exclusively end-on engagement at the N-terminus. All three loading strategies yielded single Hsp104-Cy3:β-casein-Cy5 complexes spanning the accessible FRET range (fig. S3A-C).

### ebFRET model-order comparison strongly favors three conformational states

To determine how many conformational states are required to describe Hsp104 conformational dynamics during substrate translocation, we compared ebFRET models with K = 2–6 states using the variational lower bound as approximate log evidence (the log probability of the observed data under each model, penalizing complexity through marginalization over all parameters). Across all three conditions (Hsp104^N728A^–β-casein, WT Hsp104–β-casein, and WT Hsp104–Sup35-NM fibrils), the mean lower bound per series increased monotonically with K (Table S4), but the gains were highly unequal: moving from 2 to 3 states produced by far the largest improvement of ∼10^3^ units, after which each further addition yielded progressively smaller increments (Table S5). Although these later increments still formally exceed ΔL ≳ 5, expected given the large number of observations, their small, monotonically decreasing magnitude reflects diminishing returns rather than genuine additional structure. This elbow at K = 3, together with the mixture model (Table S3), identifies three states as the most parsimonious description. A two-state description is therefore inadequate, while models with more than three states offer limited additional explanatory power. Together, these model-order comparisons establish that three states (closed, extended, and hyperextended) are the most parsimonious description consistent with the data, and that each represents a genuine, recurrent conformation rather than a noise component or an overfitting artifact.

### Model-independent analysis of Hsp104 conformational periodicity

To evaluate the possibility of periodic, sequential conformational transitions in Hsp104, we applied a model-independent analysis using the autocorrelation function (ACF) of smFRET time trajectories (as detailed in Methods) (*63*). This approach allows detection of repeated or cyclic behaviors in the FRET signal without assuming an underlying kinetic model.

The ACF plot (fig. S9A) highlights minor FRET fluctuations (expanded y-axis: –0.05 to 0.10), with the full-range inset capturing long-range dynamics. A primary correlation 1/3 decay time (see the zoomed-in-full-range insert) of ∼2.0 s, ∼8.8 s, and ∼3.2 s was observed for Hsp104^N728A^ in the presence of casein and ATP (fig. S9A), WT Hsp104 in the presence of casein and ATP:ATPγS (fig. S9B), and WT Hsp104 in the presence of Sup35-NM fibrils and ATP (fig. S9C), respectively. Power spectral density (PSD) analysis revealed a weak peak near ∼0.01 s⁻¹ (∼100 s periodicity).

To benchmark our experimental ACF results, we generated simulated smFRET traces based on three hypothetical kinetic models for Hsp104 conformational cycling:

1. A **strictly sequential motor model** (fig. S9D - a; alternative state assignments and rates in fig. S10A, B - a),
2. A **fully stochastic model** allowing all transitions (fig. S10A, B - b),
3. A **stochastic model with restricted transition pathways** (fig. S9D-b; fig. S10A,B-c).

Each model employed three FRET states (with efficiencies 0.25, 0.45, and 0.75), corresponding to hyperextended, extended, and closed conformations of the Hsp104 hexamer seam interface. Alternative FRET assignments and kinetic parameters are shown in fig. S10. Simulations were performed using the Gillespie Monte Carlo algorithm (*76, 80*), incorporating photon shot noise, trace lengths, and FRET transition rates consistent with experimental conditions. For most simulations, the transition rate was set to 0.33 s^-1^, corresponding to the ATPase rate of Hsp104^N728A^ (*k*_cat_ = 19.5 min^-1^ ≈ one ATP hydrolysis every 3.08 s) (*46*). Additional simulations were conducted with a faster transition rate of 1 s^-1^ to test frequency sensitivity (fig. S10B).

To further validate this observation of stochastic conformational change, we simulated smFRET traces under the two stochastic models. Neither the fully stochastic nor the restricted-path stochastic models produced distinct peaks in the Fourier power spectra that would correspond to 1/6 or 1/12 of the ATPase cycle time. Instead, the restricted-path model showed a low-amplitude broad peak around ∼0.01 s⁻¹ (∼100 s cycle), a timescale that aligns with weak features occasionally observed in experimental traces. However, this periodicity is too slow to be reliably resolved within the time window of our experiments and does not imply a sequential stepping mechanism.

As summarized in fig. S9 and fig. S10:

- The **sequential model** produced strong Fourier peaks at ∼0.05 s⁻¹ (20 s cycle) for 0.33 s⁻¹ rates and at ∼0.17 s⁻¹ (6 s cycle) for 1 s⁻¹ rates, corresponding to six exponentially-timed transitions per cycle.
- The **fully stochastic model** lacked dominant frequency peaks, indicating random, aperiodic transitions.
- The **stochastic model with restricted pathways** produced a broad, low-frequency peak (∼0.01 s⁻¹), aligning with experimental data but reflecting non-periodic behavior.

Collectively, the finding that experimental ACFs do not exhibit the requisite periodicity that a sequential model clearly evidences indicates that Hsp104 does not conform to a strictly sequential stepping mechanism, but rather acts as a biased stochastic motor, with non-random but non-periodic transition patterns, potentially optimized for diverse substrate interactions.

---

**Fig. S1.**
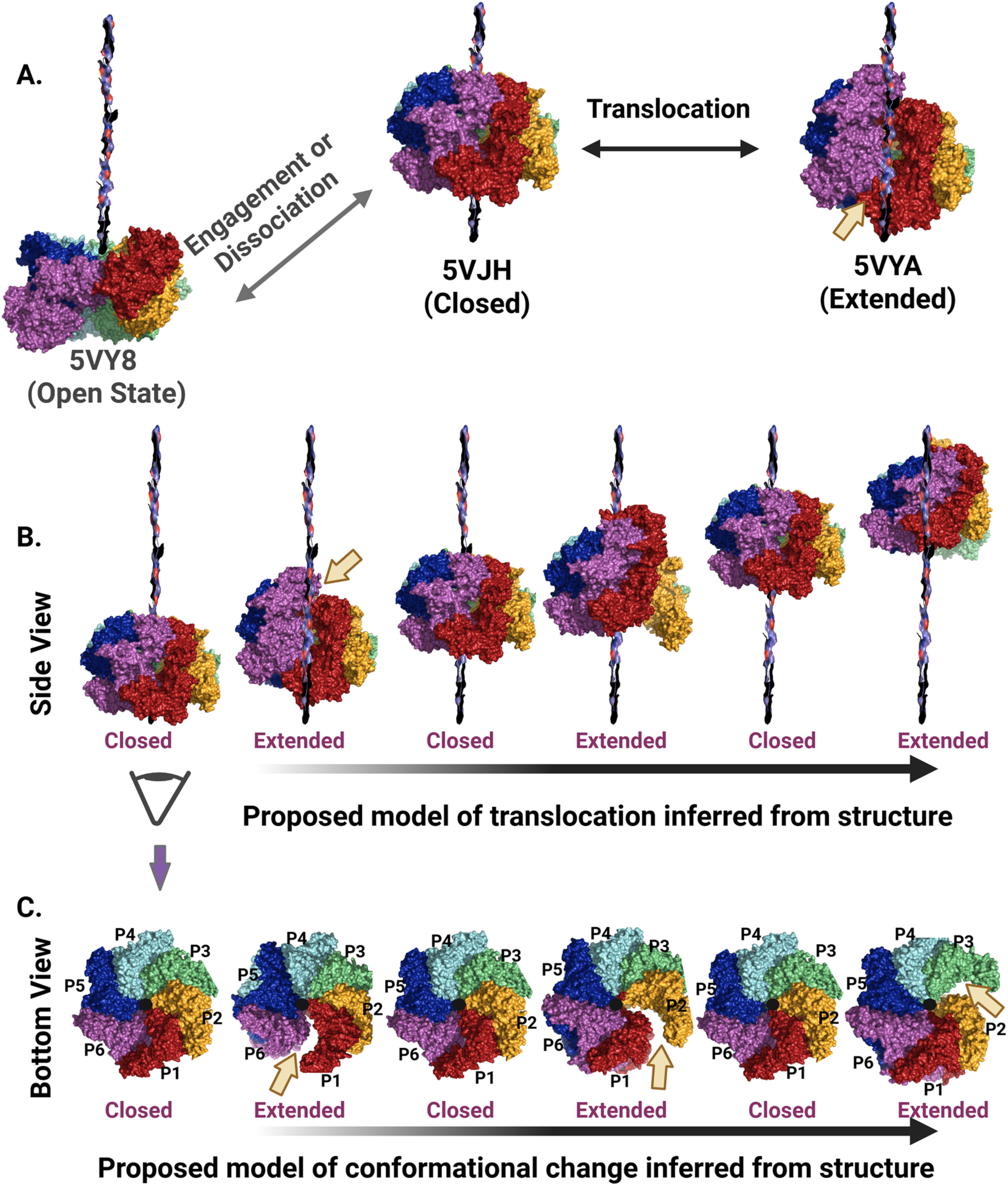
Structural model for Hsp104 translocation inferred from cryo-EM. **(A)** Cryo-EM structures of Hsp104 in open (ADP-bound; PDB 5VY8), substrate-engaged closed (PDB 5VJH), and extended (PDB 5VYA) conformations. Protomers are colored consistently across structures to track their relative positions. The threaded polypeptide is modeled as a blue–black–red rod, and the P6–P1 seam is indicated by an arrow. An additional AMP-PNP-bound open conformation (PDB 5KNE) has been reported but is not shown (*28, 33*). **(B)** Side views illustrate the previously proposed structural mechanism in which a left-handed helical, ratchet-like cycling between closed and extended states at the spiral interface drives unidirectional substrate translocation through the pore, with the seam position (arrow) marking its inferred progression (*5, 28*). **(C)** Bottom views of the same conformational series (rotated 180°) again indicating closed and extended states. Note how the seam (arrow) shifts as subunits interconvert between closed and extended conformations during this inferred translocation cycle (*5, 28*). Protomers P1-P6 are indicated.

**Fig. S2.**
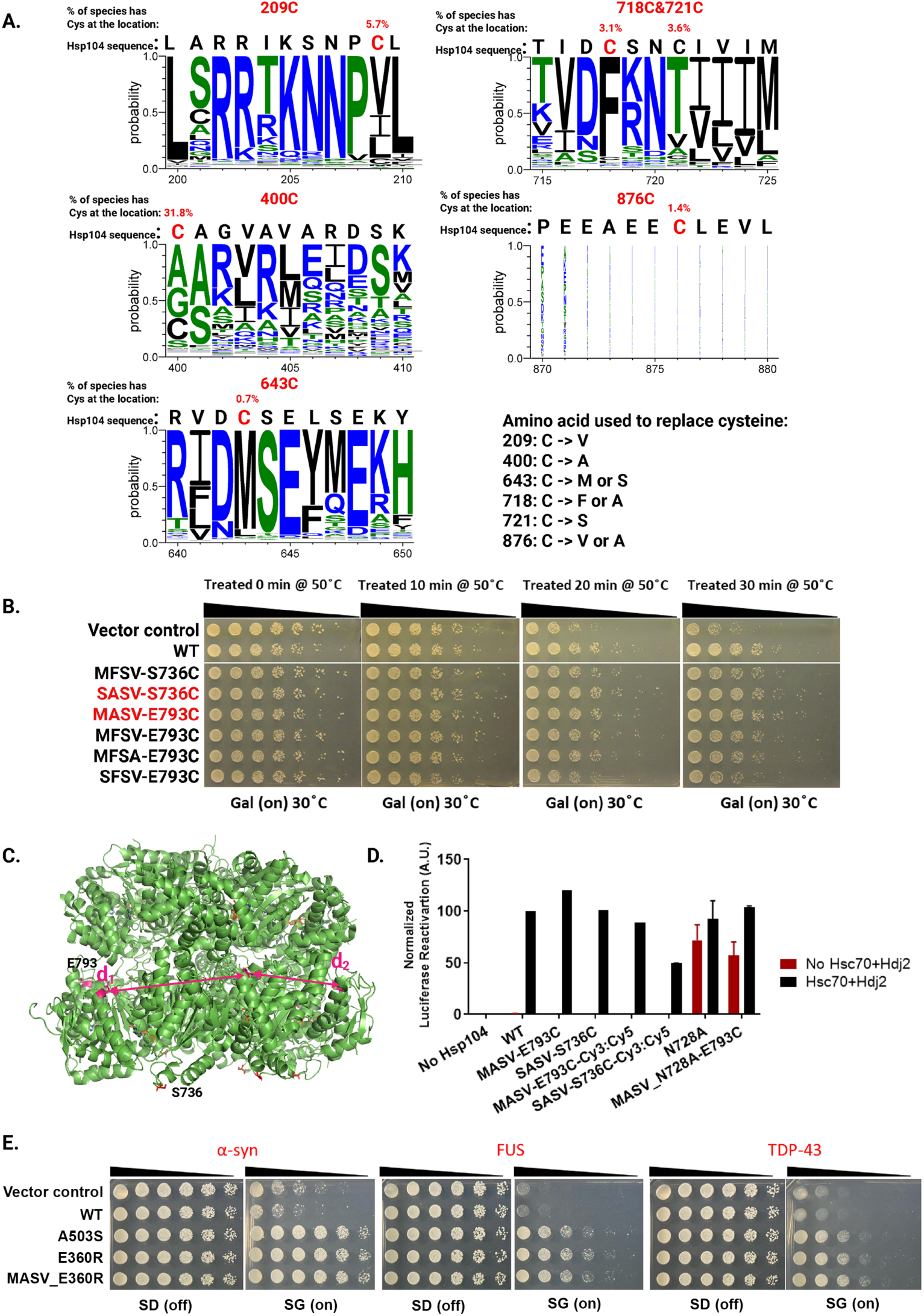
Single-cysteine variants of Hsp104 retain wild-type activity. **(A)** GREMLIN conservation analysis of the Hsp104 protein family. Sequence logos from 7,859 Hsp104 homologs show that none of the six native cysteine positions in *S. cerevisiae* Hsp104 are strongly conserved. Substitutions used to generate cysteine-light Hsp104 variants are indicated. **(B)** Thermotolerance assay assessing survival of Δ*hsp104* yeast transformed with an empty vector or the indicated Hsp104 variants. Cells were induced for 5 h at 30°C, heat-shocked at 50°C for 0, 10, 20, or 30 min, and spotted as fivefold serial dilutions on galactose-containing plates. One representative experiment is shown (n = 2). **(C)** Selection of single-cysteine labeling sites in Hsp104. Solvent-exposed residues S736 (orange/red) and E793 (magenta) were selected for site-specific labeling. The Hsp104 hexamer is shown as a green cartoon based on the closed-state cryo-EM structure (PDB 5VJH); magenta lines indicate the two intersubunit E793–E793 distances (d_1_ and d_2_). **(D)** Luciferase disaggregation and reactivation by the indicated Hsp104 variants (1 µM) in the absence (red bars) or presence (black bars) of Hsc70 (an Hsp70) (0.167 µM) and Hdj2 (an Hsp40) (0.167 µM). Values represent means ± SEM (n = 2). **(E)** Potentiated Hsp104^E360R^ retains activity in the MASV-E793C background. Δ*hsp104* yeast expressing α-synuclein–YFP, FUS, or TDP-43 from galactose-inducible promoters were transformed with the indicated Hsp104 variants. Five-fold serial dilutions were spotted onto glucose (SD; induction off) and galactose (SG; induction on) plates. Empty vector and WT Hsp104 serve as negative controls, and Hsp104^A503S^ is a positive control for potentiation. One representative experiment is shown (n = 2).

**Fig. S3.**
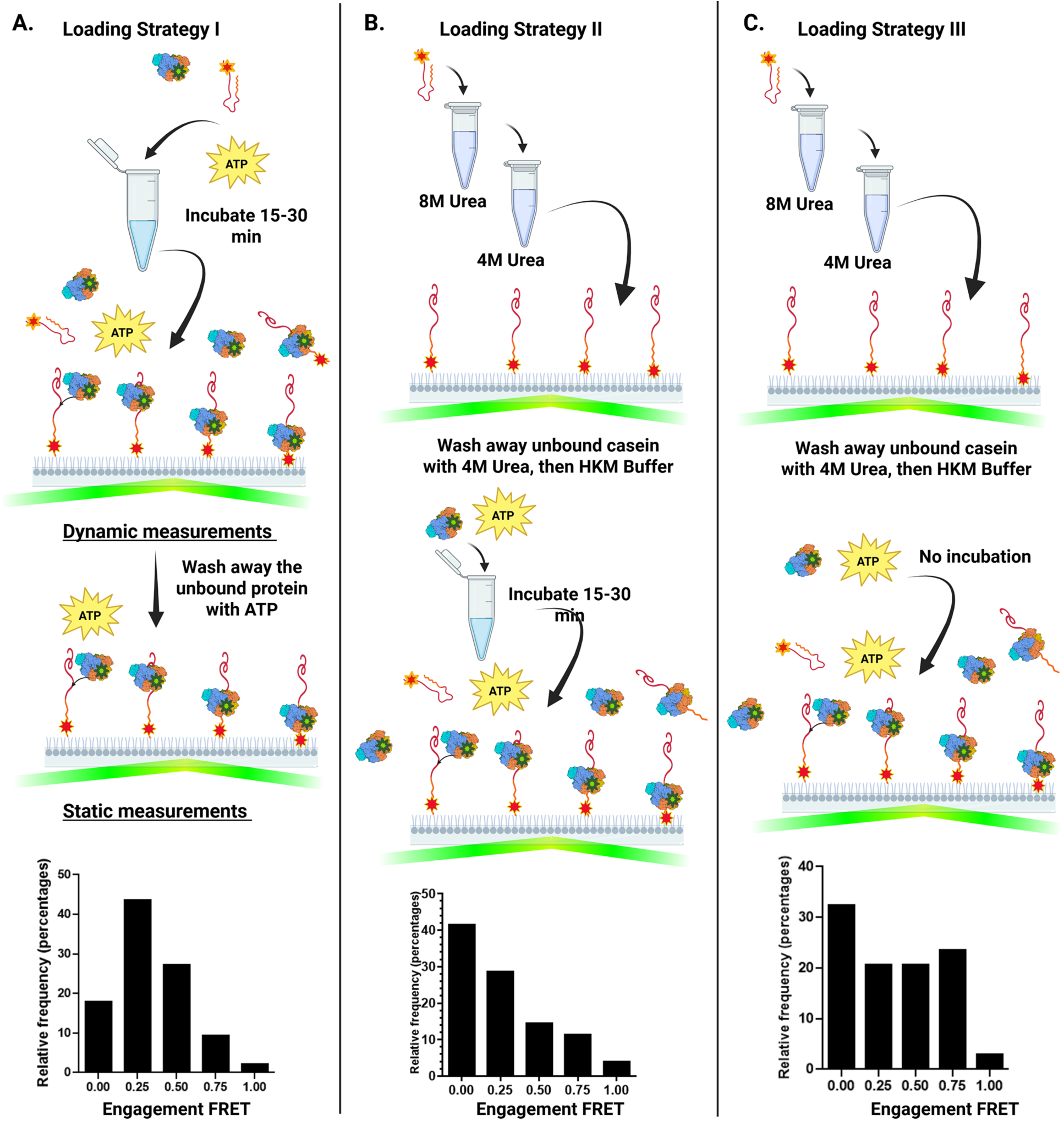
Sample preparation and loading strategies for Hsp104^N728A^ translocation experiments. **(A)** Pre-incubation strategy. Hsp104^N728A^ (120–3600 nM, monomer) and Cy5-labeled β-casein (20–600 nM) were mixed at equal concentrations in HKM150 containing 5 mM ATP and an ATP-regenerating system and incubated for 15–30 minutes at room temperature. The sample was then diluted into buffer with 5 mM ATP and illumination-buffer components to yield ∼10 nM Cy3-labeled Hsp104^N728A^ and Cy5-labeled β-casein before imaging. For dynamic measurements, unbound Hsp104^N728A^ was retained in solution and two to three 5-minute movies were acquired. For static measurements, unbound protein was removed by washing with 5 mM ATP before recording a 5-minute movie. The histogram shows FRET efficiencies at initial Hsp104^N728A^ engagement; 46.2% of hexamers engaged at a FRET efficiency of ∼0.2 (N = 184). **(B)** Pre-diluted casein strategy. Cy5-labeled β-casein (1 µM in 8 M urea) was diluted 100-fold into 4 M urea immediately before use, loaded onto the slide for 2 min, and washed sequentially with 4 M urea and HKM150 to remove unbound substrate. In parallel, Hsp104^N728A^ was pre-incubated with ATP for 15–30 minutes before loading onto the slide. The histogram shows initial engagement events; 41.6% began at zero FRET, the most frequent initial position (N = 221). **(C)** Direct-injection strategy. Cy5-labeled β-casein was prepared and immobilized as in (B). Hsp104^N728A^ and ATP were introduced directly into the imaging channel, and data acquisition began immediately after injection. One 5-minute movie was collected per injection. This strategy yielded relatively few FRET pairs per movie, with engagement events broadly distributed across the accessible FRET range (N = 34).

**Fig. S4.**
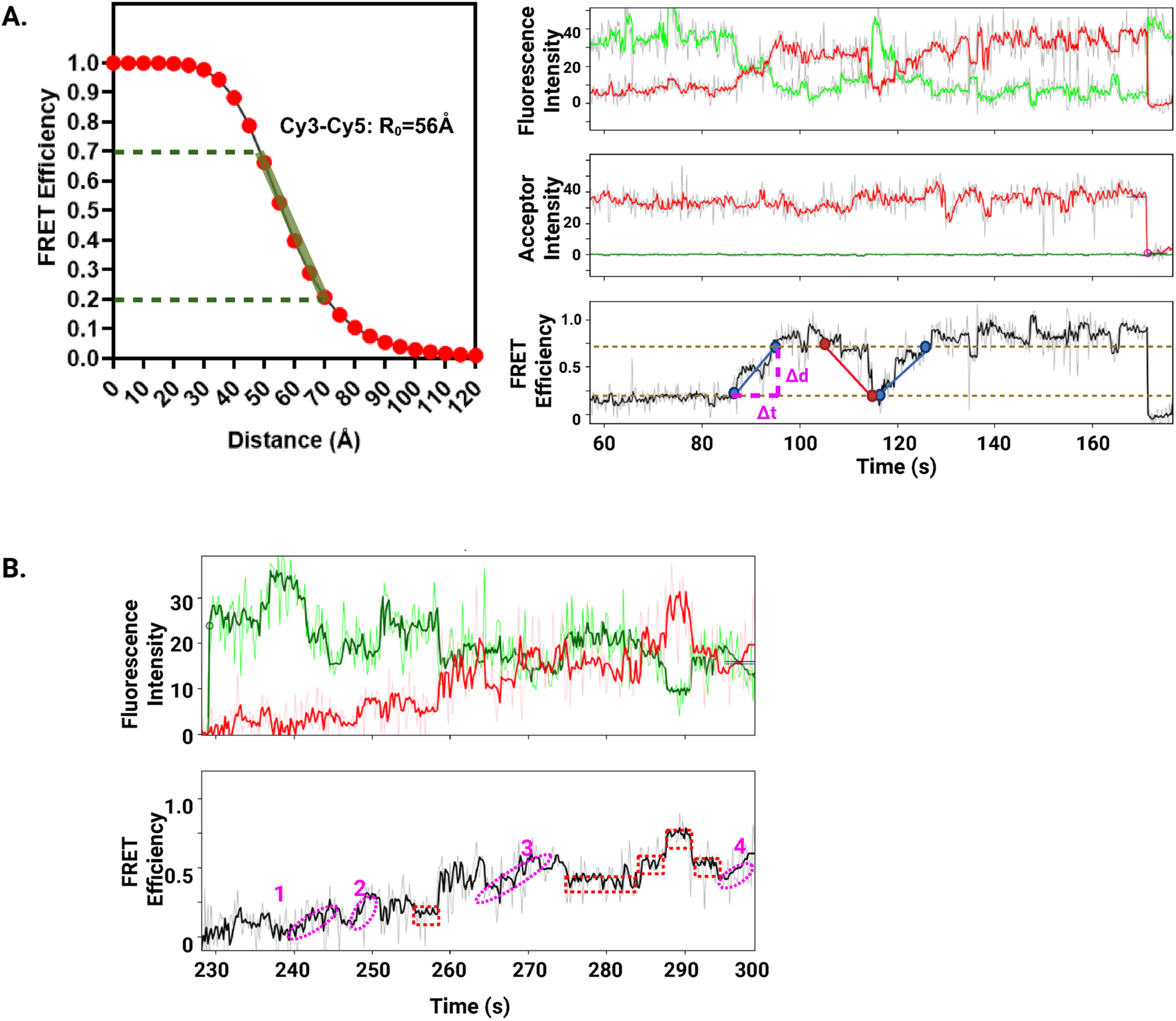
Quantification of Hsp104-Cy3 translocation events on β-casein-Cy5. **(A)** Method for estimating translocation rates from smFRET trajectories. Left: Approximate linear relationship between Cy3–Cy5 FRET efficiency and distance over the FRET range used for rate analysis (0.2–0.7; green line); dashed horizontal lines mark the boundaries of this range. Right: Representative donor (green), acceptor (red), and FRET-efficiency (black) trajectories. Three distance-change events within the defined FRET range are indicated; rates were calculated as Δd/Δt. Blue segments denote negative Δd, corresponding to motion toward the slide, and the red segment denotes positive Δd, corresponding to motion away from the slide. **(B)** Representative smFRET trajectories showing incremental distance changes (pink dashed circles) and pauses followed by rapid distance changes (red dashed boxes).

**Fig. S5.**
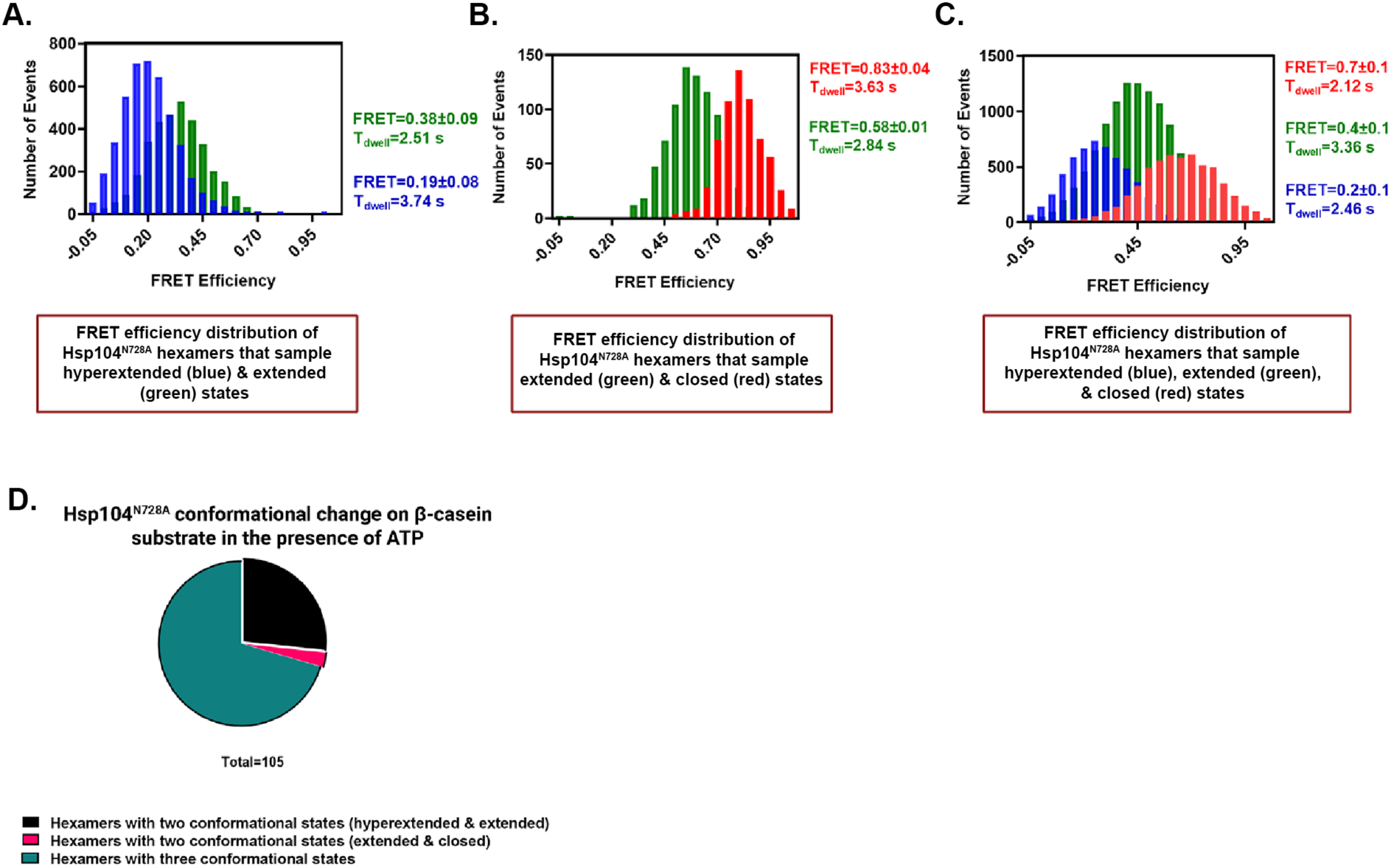
Subpopulation analysis supports two- and three-state FRET classifications of Hsp104^N728A^ hexamers on β-casein. **(A-C)** FRET-efficiency distributions for Hsp104^N728A^ trajectories on β-casein classified as sampling hyperextended and extended states (A), extended and closed states (B), or hyperextended, extended, and closed states (C). For each subpopulation, ebFRET analysis was performed with the model order fixed to the assigned category (two states in A and B; three states in C), yielding the mean FRET efficiency and modal dwell time for each resolved state. Category assignments were initialized using the per-trace Bayesian mixture model (Table S3) and refined by fixed-state ebFRET fits (Materials and Methods). Accordingly, the percentages shown reflect the ebFRET-refined assignments and differ modestly from the raw Table S3 classifications; the three-state category predominates under both approaches. **(D)** Fractions of Hsp104^N728A^ hexamers on β-casein classified as sampling two or three conformational states at labeled interprotomer interfaces. Of 105 hexamer trajectories, ∼26.7% sampled hyperextended–extended states (black), ∼2.9% sampled extended–closed states (pink), and ∼70.5% sampled all three states (teal).

**Fig. S6.**
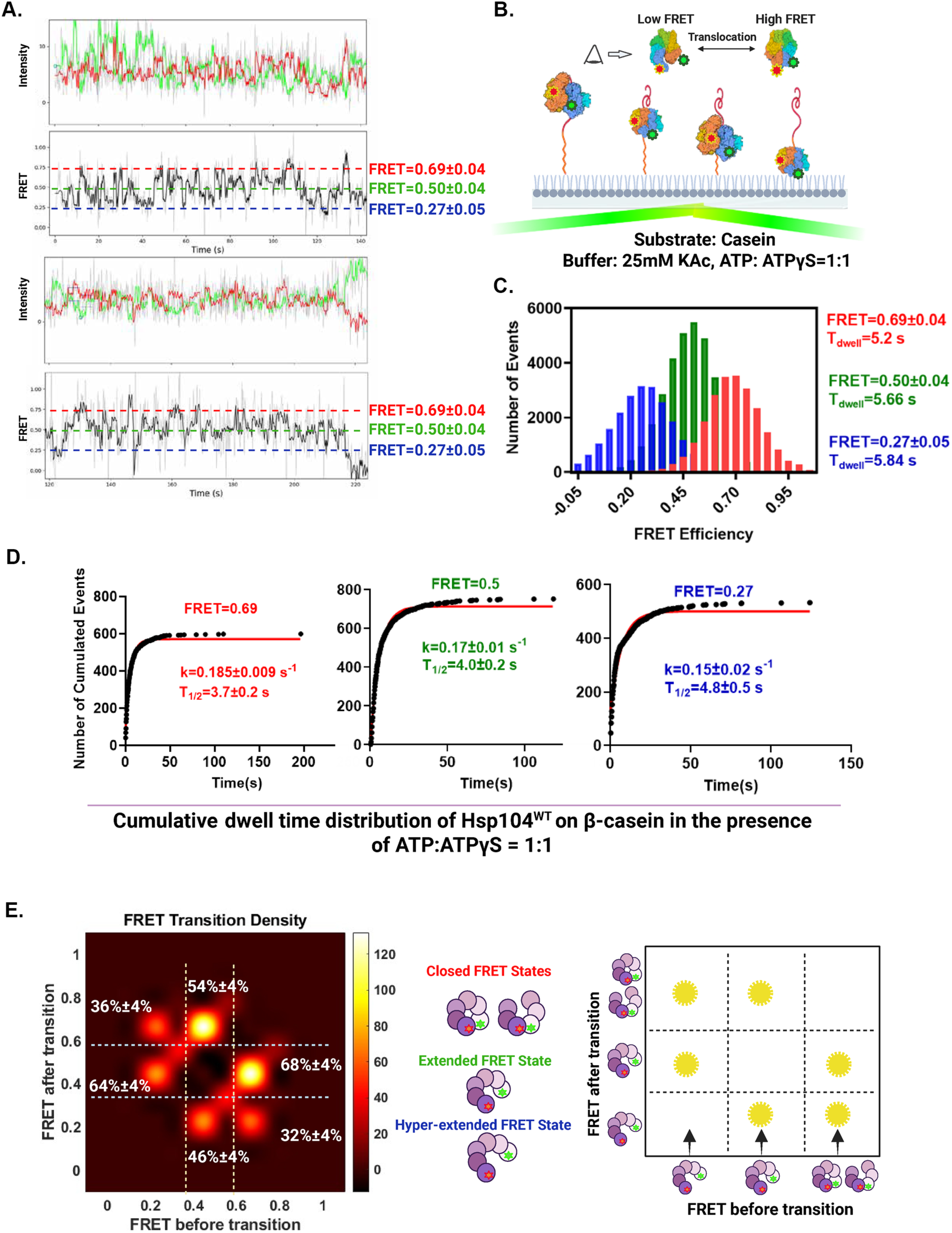
WT Hsp104 exhibits three-state, biased stochastic conformational dynamics on β-casein. **(A)** Conformational dynamics of WT Hsp104 on β-casein in ATP:ATPγS. Representative smFRET trajectories of Cy3/Cy5-labeled WT Hsp104 hexamers on β-casein in HKM25 with 2.5 mM ATP and 2.5 mM ATPγS, showing donor (green), acceptor (red), and FRET efficiency (black) with Hidden-Markov state optimization. Dashed horizontal lines indicate the mean FRET efficiency for each assigned state (closed, 0.69±0.04; extended, 0.50±0.04; hyperextended, 0.27±0.05). Across the data set (n = 149 hexamers from five independent experiments), the ebFRET-refined trajectory classification used for the subpopulation analysis in fig. S7 assigned 72.5% of Hsp104 hexamers to a three-state model and 27.5% to a two-state model; the corresponding per-trace Bayesian mixture assignment (Table S3) gave 59.7% (three-state) and 40.3% (two-state). **(B)** Schematic of the experimental setup for measuring WT Hsp104 conformational changes on β-casein. **(C)** Global ebFRET analysis of the 149 hexamers using a three-state model yielded mean FRET efficiencies of ∼0.69, ∼0.50, and ∼0.27, corresponding to closed, extended, and hyperextended conformations, with mode dwell times of 5.2, 5.66, and 5.84 s, respectively. **(D)** Cumulative dwell-time distributions for each state were fitted with single exponentials to obtain state-specific rate constants and half-times (n=599, 753, and 533 dwells for closed, extended, and hyperextended states, respectively; see Equation 4). **(E)** Transition density plot (left) summarizing the preferred conformational pathways among the three states, and corresponding schematic (right). Transitions are enriched between neighboring states but also include direct switches between closed and hyperextended conformations, consistent with biased stochastic conformational stepping. The transition-probability estimates and confidence intervals are derived from nonparametric bootstrap resampling of pooled trajectories and describe the empirical kinetic landscape at labeled interprotomer interfaces.

**Fig. S7.**
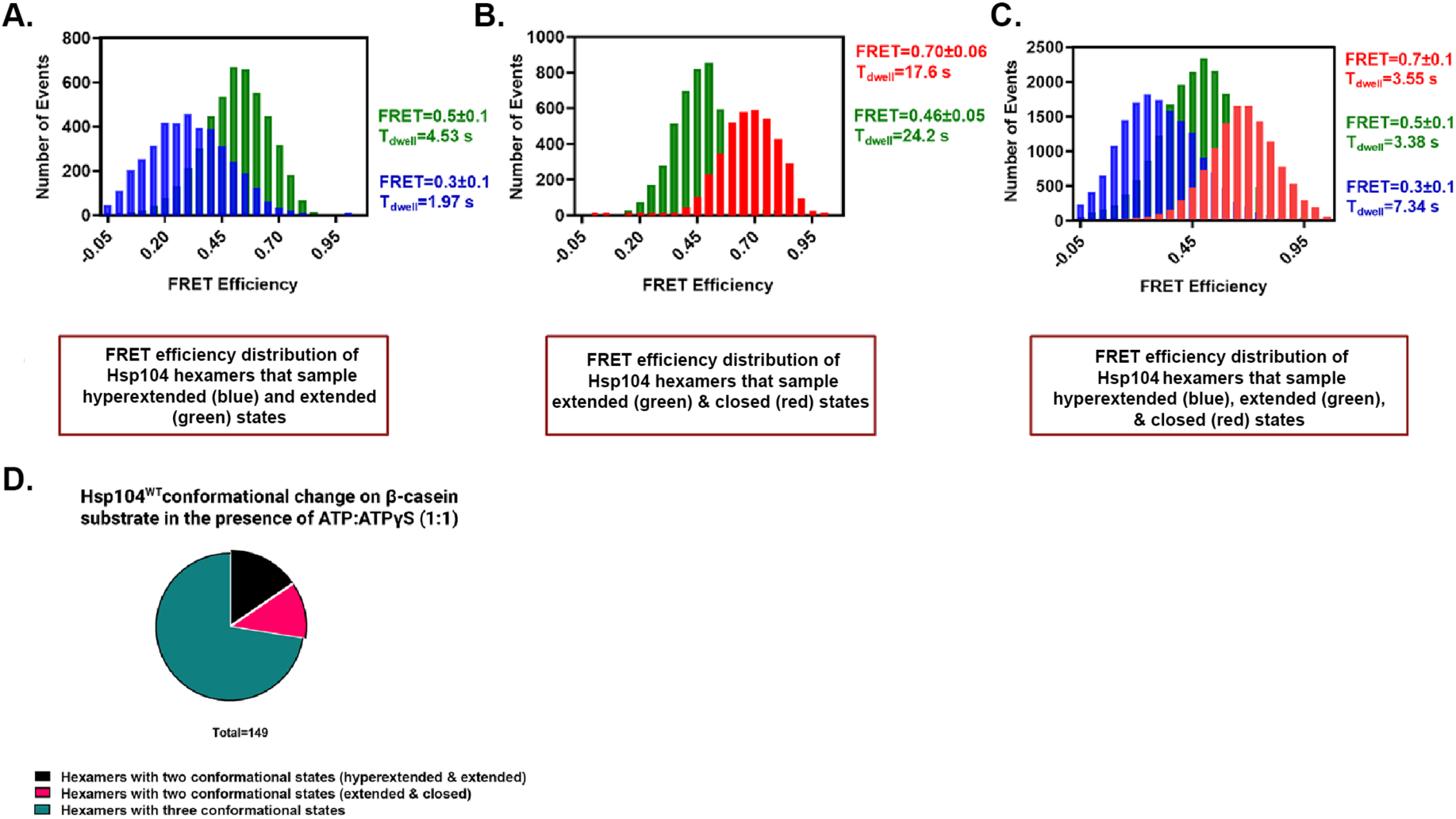
Subpopulation analysis supports two- and three-state FRET classifications of Hsp104 hexamers on β-casein. **(A-C)** FRET-efficiency distributions for Hsp104 trajectories on β-casein classified as sampling hyperextended and extended states (A), extended and closed states (B), or hyperextended, extended, and closed states (C). For each subpopulation, ebFRET analysis was performed with the model order fixed to the assigned category (two states in A and B; three states in C), yielding the mean FRET efficiency and modal dwell time for each resolved state. Category assignments were initialized using the per-trace Bayesian mixture model (Table S3) and refined by fixed-state ebFRET fits (Materials and Methods). Accordingly, the percentages shown reflect the ebFRET-refined assignments and differ modestly from the raw Table S3 classifications; the three-state category predominates under both approaches. **(D)** Fractions of Hsp104 hexamers on β-casein classified as sampling two or three conformational states at labeled interprotomer interfaces. Of 149 hexamer trajectories, ∼15.4% of hexamers sample hyperextended–extended states (black), ∼12.1% sample extended–closed states (pink), and ∼72.5% sample all three states (teal).

**Fig. S8.**
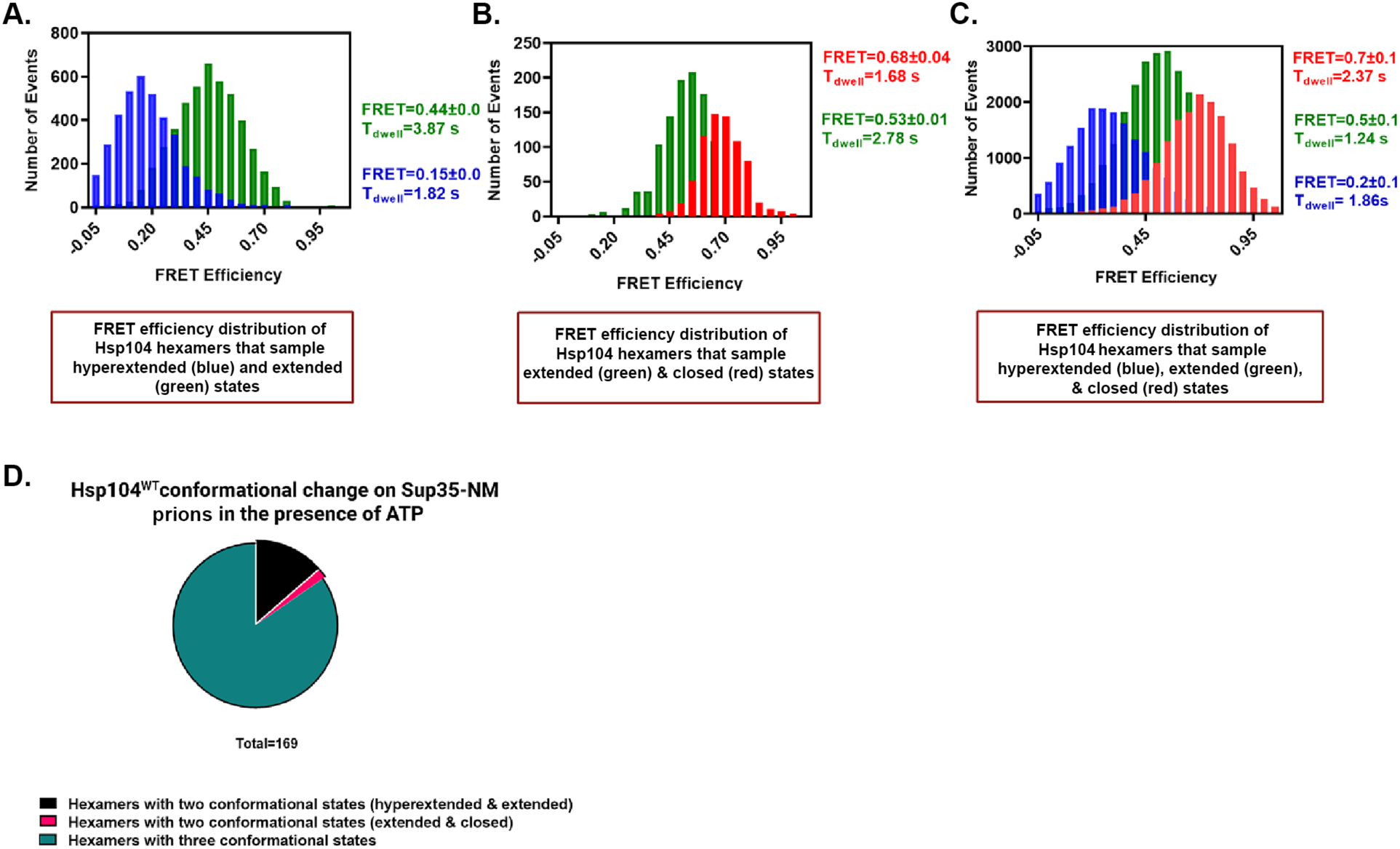
Subpopulation analysis supports two- and three-state FRET classifications of Hsp104 hexamers on Sup35-NM prions. **(A-C)** FRET-efficiency distributions for Hsp104 trajectories on Sup35-NM prions classified as sampling hyperextended and extended states (A), extended and closed states (B), or hyperextended, extended, and closed states (C). For each subpopulation, ebFRET analysis was performed with the model order fixed to the assigned category (two states in A and B; three states in C), yielding the mean FRET efficiency and modal dwell time for each resolved state. Category assignments were initialized using the per-trace Bayesian mixture model (Table S3) and refined by fixed-state ebFRET fits (Materials and Methods). Accordingly, the percentages shown reflect the ebFRET-refined assignments and differ modestly from the raw Table S3 classifications; the three-state category predominates under both approaches. **(D)** Fractions of Hsp104 hexamers on Sup35-NM prions classified as sampling two or three conformational states at labeled interprotomer interfaces. Of 169 hexamer trajectories, ∼13.6% sample hyperextended–extended states (black), ∼1.8% sample extended–closed states (pink), and ∼84.6% sample all three states (teal).

**Fig. S9.**
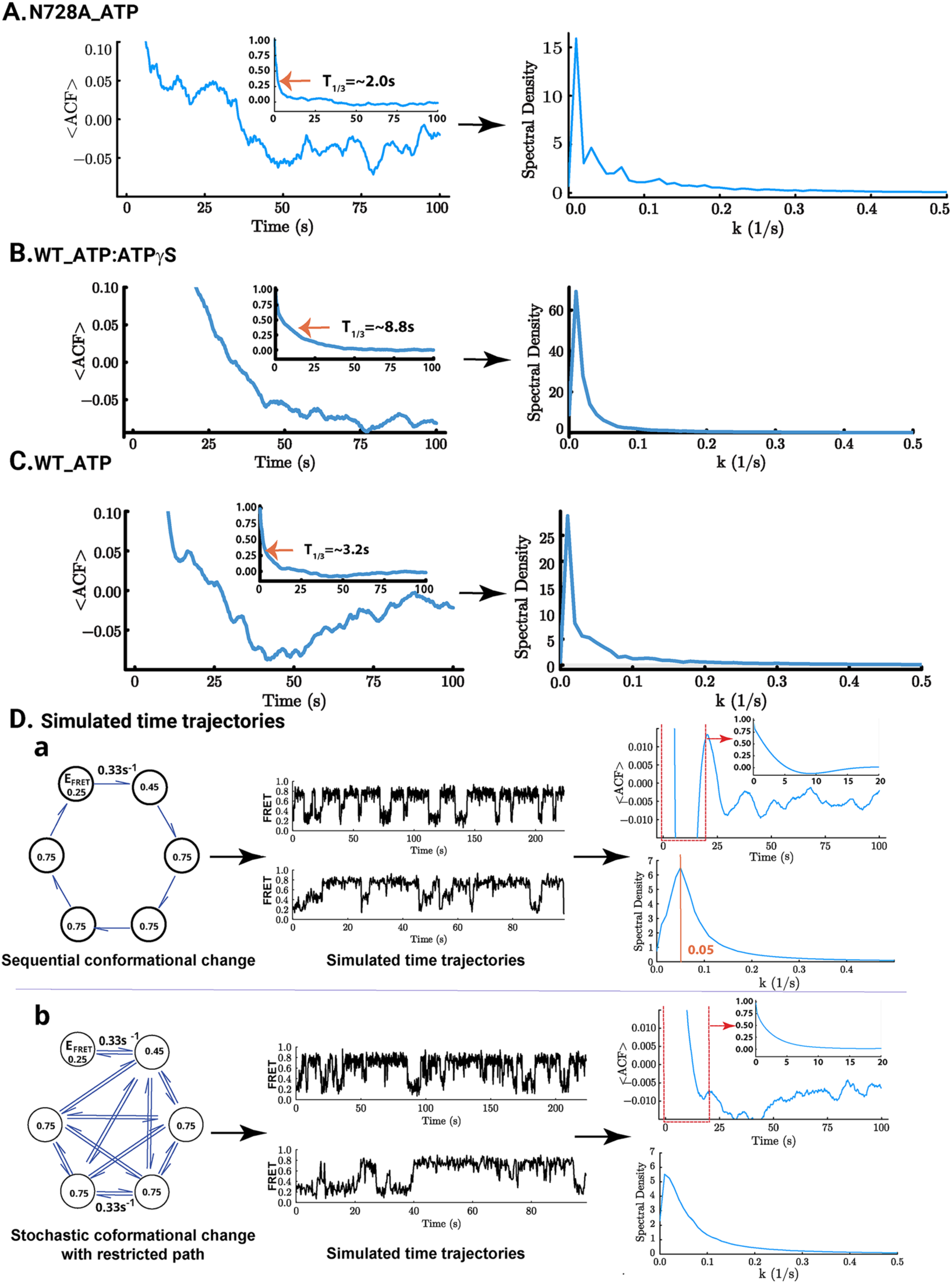
Model-independent analysis to determine the periodicity of Hsp104 conformational change. Autocorrelation function (ACF) and Fourier transform analysis of FRET time trajectories assess periodicity in Hsp104 conformational changes for (**A**) Hsp104^N728A^ bound to casein in the presence of ATP; (**B**) Hsp104 bound to casein in the presence of ATP:ATPγS; and (**C**) Hsp104 bound to Sup35-NM fibrils in the presence of ATP. Left panel: the ACF was applied to all FRET trajectories longer than 500 frames. The large plot shows an expanded view of the y-axis to highlight low-amplitude FRET fluctuations (y-axis: −0.05 to 0.10). The small plot presents a full view of the y-axis to emphasize the ACF decay time. For (**A**), the 1/3 decay time is ∼ 2.0 s. Fourier power spectral density analysis (right panel) reveals a peak at ∼0.01 s^-1^, corresponding to a ∼100 s periodicity, near the maximum observable time window. For (**B**), the 1/3 decay time is ∼ 8.8 s. On the right panel, Fourier power spectral density analysis reveals a peak at ∼0.01 s^-1^, corresponding to a ∼100 s periodicity, near the maximum observable time window. For (**C**), the 1/3 decay time is ∼3.2 s. On the right panel, Fourier power spectral density analysis reveals a peak at ∼0.01 s^-1^, corresponding to a ∼100 s periodicity, near the maximum observable time window. **(D)** Simulated FRET time traces illustrate three models of Hsp104 conformational change kinetics: **(a)** sequential motor and **(b)** stochastic motor with restricted transition pathways. FRET states (0.25, 0.45, and 0.75) are assigned to Hsp104 hexamer interfaces as shown in the left panels, with alternative assignments and transition rate constants in **fig. S10**. Simulated traces are presented in the middle panels and were generated using the Gillespie algorithm, with parameters matched to experimental conditions, including photon counts, noise, and transition rates (0.33 s^-1^), corresponding to an ATP hydrolysis rate of 19.5 min^-1^. The right panels show autocorrelation function (ACF) analysis and Fourier power spectra. The expanded view of the ACF is presented in the large plot, and a full-axis view of the ACF is plotted in the small plot. The sequential model **(a)** exhibits a periodic signal (0.05 s^-1^, ∼20 s) with a 1/3 decay time of ∼2.6 s. The calculated dwell time for 0.25, 0.45, and 0.75 FRET states are 3.0 s, 3.0 s, and 12.0 s, respectively. The restricted-path stochastic model **(b)** shows a peak at 0.01 s^-1^, which represents a ∼100 s periodicity. The 1/3 decay time is ∼2.4 s. The calculated dwell time for 0.25, 0.45, and 0.75 FRET states is 3.0 s, 0.6 s, and 3.0 s, respectively.

**Fig. S10.**
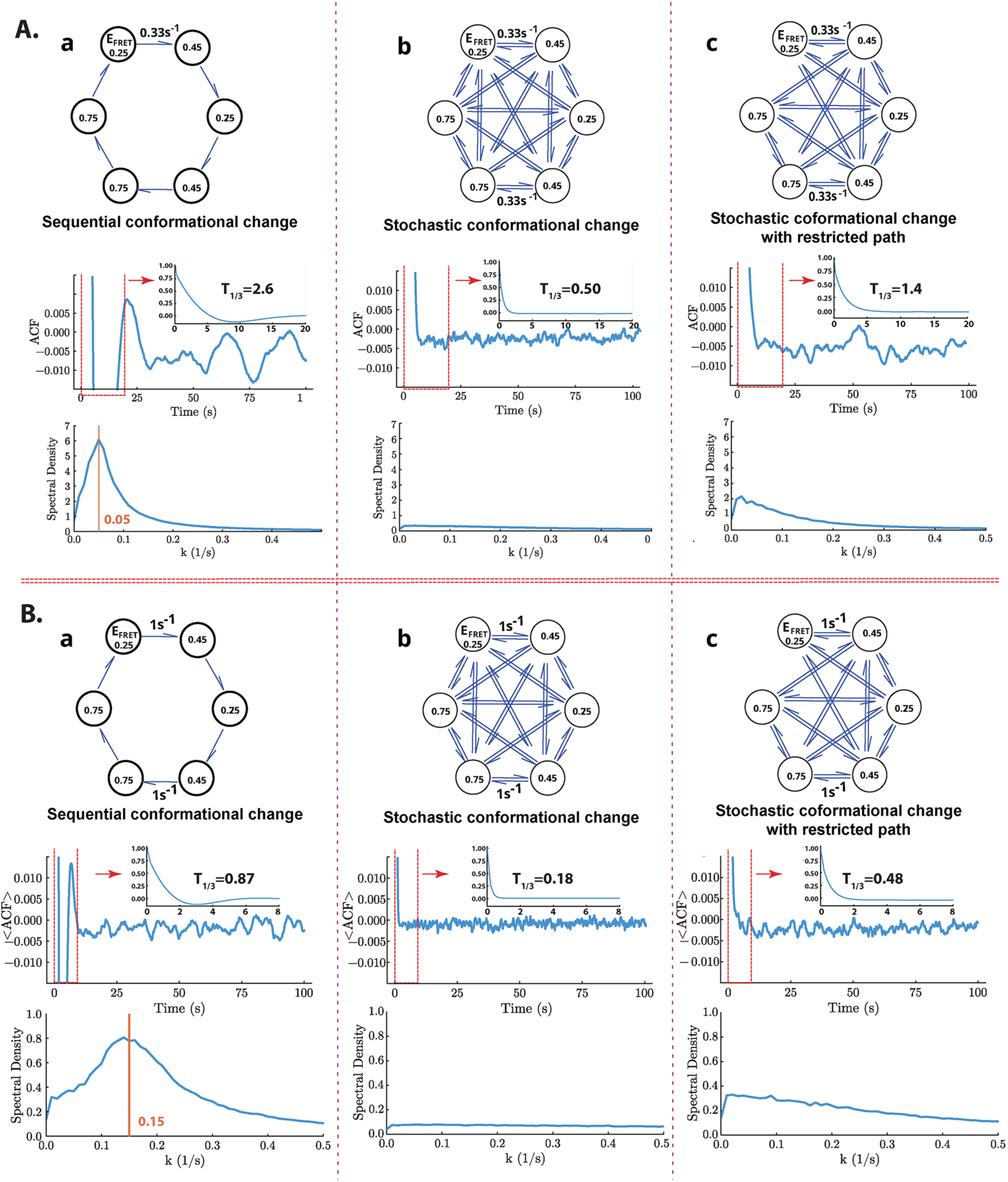
Simulations of Hsp104 conformational change with alternative FRET state assignments and transition kinetics. FRET states (0.25, 0.45, and 0.75) are assigned to Hsp104 hexamer interfaces, as illustrated in the top panels, with two sets of transition rate constants: **(A)** 0.33 s^-1^ and **(B)** 1 s^-1^. Three models of motor activity are presented: (a) sequential, (b) fully stochastic, and (c) stochastic with restricted transition pathways. The bottom panels display autocorrelation function (ACF) analyses and corresponding Fourier transform power spectra to detect the periodicity. An expanded view of the x-axis of ACF is shown in the large plot; a full axis view of the entire y-axis is shown in the inset, and the 1/3 decay time is indicated on each plot. In the sequential model **(a),** periodicity is observed: ∼20 s for the 0.33 s⁻¹ condition **(A)** and ∼6 s for the 1 s⁻¹ condition **(B)**. Calculated dwell times for the 0.25, 0.45, and 0.75 FRET states are 3.0 s, 3.0 s, and 6.0 s for **A-(a),** and 1.0 s, 1.0 s, and 2.0 s for **B-(a),** respectively. The fully stochastic model **(b)** shows no periodicity; dwell times are uniform across states at 0.75 s for all FRET states in **A-(b)** and 0.25 s for **B-(b).** The restricted-path stochastic model **(c)** exhibits a broad peak in the power spectrum at ∼0.01 s⁻¹ (∼100 s periodicity), independent of the transition rate. Dwell times for each FRET state in **A-(c)** are 1.5 s (*E*=0.25), 0.75 s (*E*=0.45), and 0.75 s (*E*=0.75), and for **B-(c),** 0.5 s (*E*=0.25), 0.25 s (*E*=0.45), and 0.25 s (*E*=0.75).

**Table S1.** Summary of Hsp104 translocation and unfolding rates.

|  | **Hsp104<sup>N728A</sup>** | **Hsp104** |
| --- | --- | --- |
| **Rate of Translocation/Unfolding**<br>**(Stopped-flow)** | 1.35 ± 0.04 aa s<sup>-1</sup> | 0.44 ± 0.02 aa s<sup>-1</sup> |
| **Rate of Translocation/unfolding**<br>**(smTIRF)** | 4.3 Å/s (4.0 Å/s - 4.6 Å/s) | 2.2 Å/s (1.8 Å/s - 2.5 Å/s), in HKM25 buffer<br>2.2 Å/s (1.9 Å/s - 2.6 Å/s), in HKM150 buffer |
| **Rate of Translocation/unfolding**<br>**(smTIRF)** | 1.26 aa/s (1.18 aa/s – 1.35 aa/s) | 0.65 aa/s (0.53 aa/s – 0.74 aa/s), in HKM25 buffer<br>0.65 aa/s (0.56 aa/s – 0.76 aa/s), in HKM150 buffer |
| **Excluded length (Stopped-flow)** | 58.6 ± 7.6 aa | 66.2 ± 12.8 aa |
| **Rate of unfolding due to ATPγS**<br>**(Stopped-flow)** | 0.07 ± 0.02 aa s<sup>-1</sup> | 0.12 ± 0.03 aa s<sup>-1</sup> |

Translocation and unfolding rates for Hsp104<sup>N728A</sup> and WT Hsp104 were determined by smTIRF and stopped-flow assays. Rates are reported in amino acids per second (aa/s) or angstroms per second (Å/s), as indicated. Single-molecule rates are reported with 95% confidence intervals. Stopped-flow rates are reported as slopes from linear regression, with errors of the mean from three independent replicates for each experimental condition. “Excluded length” denotes the non-translocated segment inferred from the y-intercept of plots of peak time versus substrate length. ATPγS-dependent unfolding rates report background activity measured in the absence of ATP.

**Table S2.** Calculated FRET efficiencies based on inter-protomer distances measured from cryo-EM structures.

| Hexamer conformation | PDB | P1-P2 | P2-P3 | P3-P4 | P4-P5 | P5-P6 | P6-P1 |
| --- | --- | --- | --- | --- | --- | --- | --- |
| ATPγS-Extended | 5VYA | 0.73 | 0.74 | 0.72 | 0.76 | 0.76 | 0.41 |
| ATPγS-Closed | 5VJH | 0.72 | 0.70 | 0.75 | 0.75 | 0.77 | 0.64 |
| ADP-Open | 5VY8 | 0.68 | 0.68 | 0.63 | 0.64 | 0.68 | 0.04 |
| AMP-PNP-Open | 5KNE | 0.68 | 0.70 | 0.66 | 0.73 | 0.74 | 0.04 |

Inter-protomer distances between E793 residues of adjacent protomers were measured in the indicated cryo-EM structures (see fig. S2C) and used to calculate FRET efficiencies using Equation 1 (Methods). Two distinct ATPγS-bound Hsp104 conformations are resolved (highlighted in blue): an extended state (PDB 5VYA) and a closed state (PDB 5VJH). FRET efficiencies at the P6–P1 seam interface (red) are markedly lower in the ADP- and AMP-PNP-bound open states than in the ATPγS-bound closed and extended states. Within the ATPγS-bound states, the P6–P1 FRET efficiency is substantially lower in the extended state (0.41) than in the closed state (0.64), consistent with widening of the seam during the closed-to-extended conformational transition.

**Table S3.** Bayesian mixture model analysis of Hsp104 conformational states identified by smFRET.

| Experiment | # of data sets | E<sub>1</sub> | E<sub>2</sub> | E<sub>3</sub> |
| --- | --- | --- | --- | --- |
| N728A_Casein 2 states (1→2) | 32 | 0.24 ± 0.05 | 0.47 ± 0.05 |  |
| N728A_Casein 2 states (2→3) | 6 |  | 0.52 ± 0.08 | 0.73 ± 0.07 |
| N728A_Casein 3 states | 67 | 0.23 ± 0.07 | 0.48 ± 0.08 | 0.74 ± 0.09 |
| WT_Casein_ATP_ATPγS 2 states (1→2) | 43 | 0.27 ± 0.08 | 0.52 ± 0.07 |  |
| WT_Casein_ATP_ATPγS 2 states (2→3) | 17 |  | 0.44 ± 0.05 | 0.69 ± 0.05 |
| WT_Casein_ATP_ATPγS 3 states | 89 | 0.22 ± 0.07 | 0.48 ± 0.07 | 0.74 ± 0.08 |
| WT_Sup35-NM fibrils 2 states (1→2) | 27 | 0.19 ± 0.06 | 0.48 ± 0.07 |  |
| WT_Sup35-NM fibrils 2 states (2→3) | 6 |  | 0.40 ± 0.06 | 0.68 ± 0.09 |
| WT_Sup35-NM fibrils 3 states | 136 | 0.20 ± 0.06 | 0.48 ± 0.06 | 0.74 ± 0.08 |

**Table S4.** Mean ebFRET variational lower bounds for model orders K = 2–6 across Hsp104–substrate conditions.

| Condition | 2 states<br>(K=2) | 3 states<br>(K=3) | 4 states<br>(K=4) | 5 states<br>(K=5) | 6 states<br>(K=6) |
| --- | --- | --- | --- | --- | --- |
| N728A-β-casein | $2.449 \times 10^3$ | $3.065 \times 10^3$ | $3.292 \times 10^3$ | $3.446 \times 10^3$ | $3.581 \times 10^3$ |
| WT-β-casein | $6.941 \times 10^3$ | $8.797 \times 10^3$ | $9.549 \times 10^3$ | $9.972 \times 10^3$ | $1.031 \times 10^4$ |
| WT-Sup35-NM<br>fibrils | $3.467 \times 10^3$ | $4.488 \times 10^3$ | $4.893 \times 10^3$ | $5.109 \times 10^3$ | $5.330 \times 10^3$ |

**Table S5.**
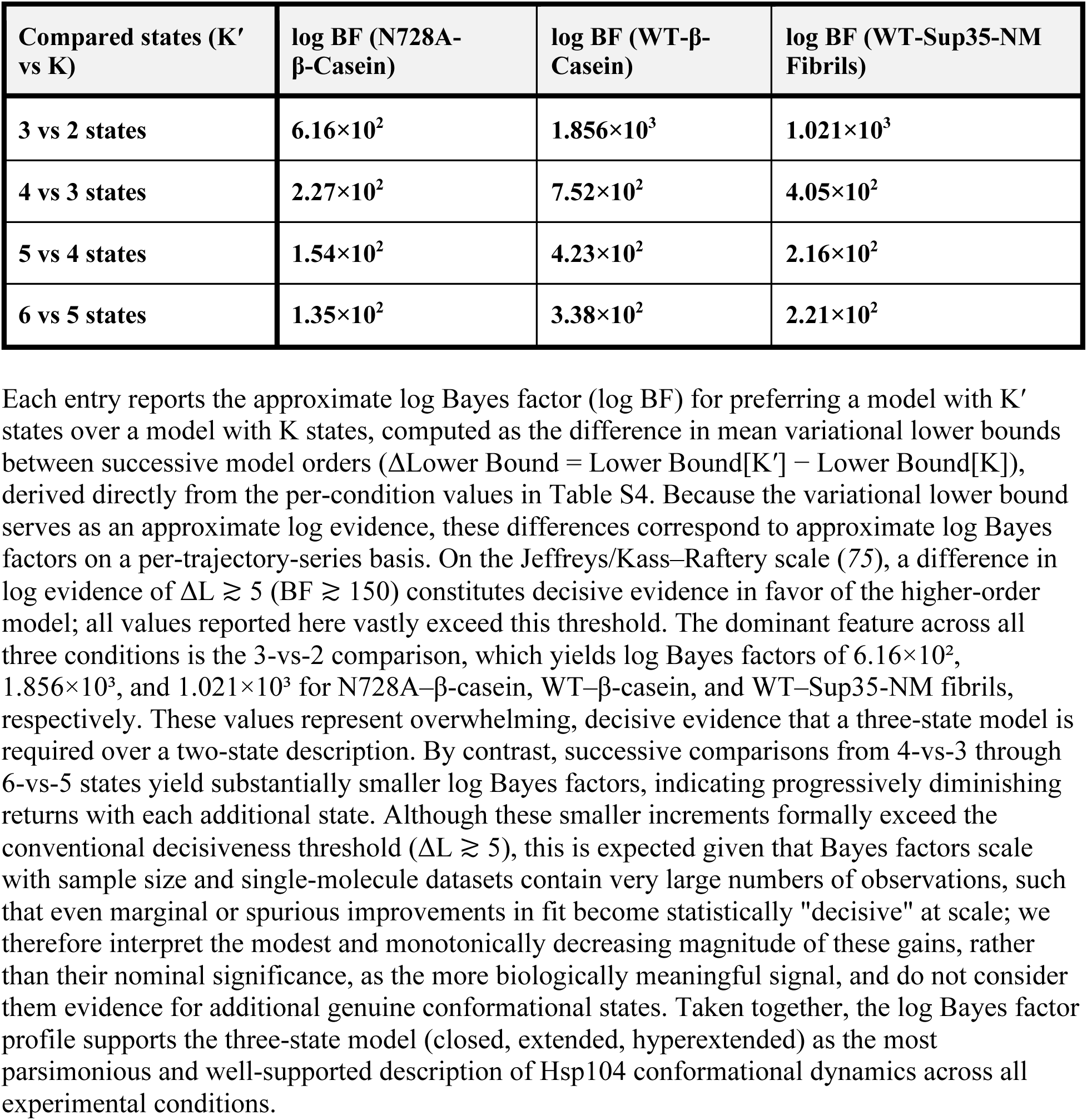
Approximate log Bayes factors comparing successive ebFRET model orders for Hsp104 conformational state fits.

| Compared states (K' vs K) | log BF (N728A-β-Casein) | log BF (WT-β-Casein) | log BF (WT-Sup35-NM Fibrils) |
| --- | --- | --- | --- |
| 3 vs 2 states | $6.16 \times 10^2$ | $1.856 \times 10^3$ | $1.021 \times 10^3$ |
| 4 vs 3 states | $2.27 \times 10^2$ | $7.52 \times 10^2$ | $4.05 \times 10^2$ |
| 5 vs 4 states | $1.54 \times 10^2$ | $4.23 \times 10^2$ | $2.16 \times 10^2$ |
| 6 vs 5 states | $1.35 \times 10^2$ | $3.38 \times 10^2$ | $2.21 \times 10^2$ |

**Table S6.** Mean fractional occupancy of closed, extended, and hyperextended conformational states across experimental conditions.

| Condition | FRET Values<br>(State 1 / 2 / 3) | Hyperextended<br>(State 1) | Extended<br>(State 2) | Closed<br>(State 3) | Combined<br>(State 1 + 2) |
| --- | --- | --- | --- | --- | --- |
| N728A-β-casein | 0.23 / 0.45 / 0.74 | 39.9% ± 2.7% | 38.4% ± 1.9% | 21.8% ± 2.3% | 78.3% |
| WT-β-casein | 0.27 / 0.50 / 0.69 | 28.8% ± 2.9% | 42.6% ± 0.1% | 28.6% ± 2.5% | 71.4% |
| WT-Sup35-NM<br>fibrils | 0.18 / 0.43 / 0.65 | 26.6% ± 2.5% | 41.5% ± 1.1% | 31.9% ± 2.7% | 68.1% |
| Rotary null | — | ~8.3% | ~8.3% | ~83.3% | ~16.7% |

**Table S7.** Bootstrap-estimated Hsp104 three-state transition probabilities (%, 95% CI) for N728A–β-casein, WT–β-casein, and WT–Sup35-NM fibril conditions.

| | Hsp104 three-state transition probabilities $P_{ij}$ (95% CI) | | |
| --- | --- | --- | --- |
| *Transition (from → to)* | *N728A-β-casein $P_{ij}$* | *WT-β-casein (ATP/ATPγS) $P_{ij}$* | *WT-Sup35-NM fibrils $P_{ij}$* |
| *Hyperextended → Extended (1→2)* | **61.2%** (56.3–66.0%) | **64.4%** (60.3–68.4%) | **58.6%** (55.6–61.6%) |
| *Hyperextended → Closed (1→3)* | **38.8%** (34.0–43.7%) | **35.6%** (31.6–39.7%) | **41.4%** (38.4–44.4%) |
| *Extended → Hyperextended (2→1)* | **53.9%** (49.2–58.6%) | **45.7%** (42.1–49.3%) | **49.6%** (46.9–52.5%) |
| *Extended → Closed (2→3)* | **46.1%** (41.4–50.8%) | **54.3%** (50.7–57.9%) | **50.4%** (47.5–53.1%) |
| *Closed → Hyperextended (3→1)* | **41.9%** (36.7–47.1%) | **31.7%** (28.0–35.5%) | **41.8%** (38.8–44.8%) |
| *Closed → Extended (3→2)* | **58.1%** (52.9–63.3%) | **68.3%** (64.5–72.0%) | **58.2%** (55.2–61.2%) |

Transition probabilities among the three conformational states (hyperextended, extended, and closed) were estimated from ebFRET-idealized smFRET trajectories using a nonparametric bootstrap procedure (B = 5,000 replicates; see Methods). For each condition, individual non-self transitions were mapped to one of six directed edges based on assignment to the nearest condition-specific FRET state center, and a 3 × 3 transition count matrix was row-normalized to yield empirical transition probabilities. Bootstrap confidence intervals (95% CI) were obtained using the percentile method (2.5th–97.5th percentiles of the bootstrap distribution). States 1, 2, and 3 correspond to the hyperextended (low FRET), extended (mid FRET), and closed (high FRET) conformations, respectively. Three experimental conditions are shown: Hsp104<sup>N728A</sup> on β-casein in ATP (N728A-β-casein), WT Hsp104 on β-casein in ATP/ATPγS (WT-β-casein), and WT Hsp104 on Sup35-NM prion fibrils in ATP (WT-Sup35-NM fibrils).

